# Gene activation and repression by the glucocorticoid receptor are mediated by sequestering Ep300 and two modes of chromatin binding

**DOI:** 10.1101/764480

**Authors:** Avital Sarusi Portuguez, Ivana Grbesa, Moran Tal, Rachel Deitch, Dana Raz, Ran Weismann, Michal Schwartz, Olga Loza, Myong-Hee Sung, Tommy Kaplan, Ofir Hakim

## Abstract

The transcription factor glucocorticoid receptor (GR) is a key mediator of stress response and a broad range of physiological processes. How can GR rapidly activate the expression of some genes while repress others, remains an open question due to the challenge to associate GR binding sites (GBSs) to their distant gene targets. Mapping the full 3D scope of GR-responsive promoters using high-resolution 4C-seq unravelled spatial separation between chromatin interaction networks of GR-activated and repressed genes. Analysing GR binding sites and other regulatory loci in their functional 3D context revealed that GR sequesters the co-activator Ep300 from active non-GBS enhancers in both activated and repressed gene compartments. While this is sufficient for rapid gene repression, gene activation is countered by productive recruitment of Ep300 to GBS. Importantly, in GR-activated compartments Klf4 binding at non-GBS regulatory elements cluster in 3D with GBS and antagonizes GR activation. In addition, we revealed ROR and Rev-erb transcription factors as novel co-regulators for GR-mediated gene expression.

## INTRODUCTION

Cells continuously respond to hormones, such as glucocorticoids (GC), by rapidly remodeling their transcriptional programs. GC signal through binding to the glucocorticoid receptor (GR, encoded by Nr3c1), a transcription factor of the nuclear receptor superfamily. GR binding to DNA is highly cell type-specific and relies on the interplay of GR with other transcription factors. In mouse mammary epithelial (3134) cells, activator protein 1 (AP-1) dictates a large proportion (50%) of GR binding sites (GBSs) (1) by priming regulatory elements that define the cell type-specific locations of accessible chromatin for subsequent GR binding (2, 3). In addition, GR can utilize these accessible sites to interact with other factors to induce variable transcription patterns (4–6). While significant progress has been made in understanding how these regulatory sites are initiated and maintained, less is known about the mechanisms that determine the specific function of GR sites in activating or repressing transcription. Gene activation is thought to be primarily controlled by direct binding of GR to its recognition element (GRE) and subsequent recruitment of co-activators such as SRC-1, GRIP1 and CBP/EP300 (7–10). Models for transcriptional repression include direct and indirect GR binding to DNA. One of the proposed mechanisms, trans-repression, stems from the indirect binding of GR to regulatory loci lacking GR recognition elements (GREs). In this case, tethering of GR to DNA-bound transcription factors like AP-1 and nuclear factor-kB (NF-κB) disrupts their association with co-activators and thereby antagonizes their activity (10–13). However genome-wide analyses indicate that the association of AP-1 and NF-κB on GR binding sites is common to both up- and down-regulated genes, and thus cannot explain the opposite transcriptional responses (1, 6, 14).

Gene repression by direct GR binding to DNA was proposed to be mediated by competition and steric hindrance to binding of other transcription factors with overlapping binding sites (9). In several cases, direct GR binding to “negative” GR elements (nGRE) can recruit the NCoR1-HDAC3 repressing complex (15–17). However, nGRE motifs or footprints are found at GR binding sites to a similar extent near activated and repressed genes (14, 18). This suggests that the GR recognition element (GRE or nGRE) alone is insufficient to globally classify the opposite transcriptional effects of GR. Another mechanism of transcriptional repression by GR involves competition with other transcription factors for a limited amount of co-activators (19). However, variation in the cellular levels of CBP/EP300 was not consistent in affecting GR-mediated repression in reporter assays (20). It is possible that this and other controversies remain due to the limited number of genes studied by reporter assays, which do not reveal the complexity of multiple regulatory elements in the chromatin context.

Perhaps one of the most significant barriers for mechanistic understanding of GR-mediated regulation is the difficulty in assigning specific GBSs to their distant gene targets. Genome-wide analyses have demonstrated that the majority (∼90%) of GBSs are distant from the transcription start sites (TSS) of their nearest gene, and that GR binding events greatly exceed the number of GR-responsive genes (2, 21), suggesting that multiple GBSs and genes associate together by chromatin folding. To advance our mechanistic understanding of GR-mediated regulation it is essential to assign each GR-regulated gene to its respective distant GR regulatory elements.

Indeed, the spatial organization of genomic information is a major regulatory component of gene transcription. In recent years, it has been appreciated that the mouse (and human) chromosomes are partitioned into autonomous units of high spatial connectivity (22). These Topologically Associating Domains (TADs) are large (ranging from tens of kb to 3 Mb) chromosomal units encompassing multiple genes and regulatory elements (23). Although the borders between TADs are mostly considered cell type-invariant, chromosomal contacts between GR binding sites and their target genes (within TADs) show cell type-specificity (24). Moreover, changes in gene expression within TADs are highly correlated during differentiation and in response to external stimuli, suggesting that TADs confine the activity of regulatory elements to specific genes (25–27). However, the spatial constraints of GR sites towards their regulated genes are yet to be fully described.

To directly determine the scope of regulatory sites associated via chromosomal looping with GR-responsive genes, we applied high-throughput and high-resolution gene-centric circular chromosome conformation capture (4C-seq) for 40 GR-regulated genes. We found that GR-responsive genes cluster in 3D with multiple GBS and other regulatory elements in hormone- and transcription-independent manners. Importantly, these spatial clusters are specialized for either transcriptional activation or repression by GR. To explore features of GR regulatory polarity, we compiled catalogs of regulatory elements associated with GR-induced or -repressed genes and combined these data with multiple ChIP-seq profiles in resting and hormone-treated cells. Our results revealed that co-activator sequestering is a major mechanism for rapid transcriptional repression. Exploring the positive enhancers identified the transcription factors Klf4 and Rev-Erb/ROR as novel GR co-regulators. Interestingly, Klf4 is associated primarily with non-GBS enhancers, indicating that the spatial convergence of multiple GBS and non-GBS elements underlie the mechanistic complexity of the bipolar transcriptional response to hormone-activated GR. Overall, our results demonstrate that precise segmentation the genome according to structure and specific transcriptional responses is a powerful discovery tool to decode the complexity of transcription factor activity.

## MATERIAL AND METHODS

### Cell growth

Mouse mammary epithelial adenocarcinoma cells (3134) (28) were maintained in Dulbecco’s modified Eagle’s medium (DMEM, Biological Industries) supplemented with 10% fetal bovine serum (FBS, Biological Industries), 2 mM L-glutamine, 0.5 mg/mL penicillin-streptomycin and 1 mM sodium pyruvate (Biological Industries) in a humidified 37°C incubator with 5% CO_2_. Before treatment (EtOH or 100 nM Dexamethasone for 1 h, Sigma), cells were cultured in media containing 10% charcoal-stripped serum (CSS) (Sigma, USA). HepG2 cells (ATCC HB-8065) were cultured in DMEM media supplemented with 10% FBS, 1% penicillin-streptomycin and 1% L-glutamine.

### RNA extraction and quantification by qPCR

RNA was extracted and on-column treated with DNase I with Quick-RNA Mini Prep kit (Zymo Research). Total RNA concentration and purity were measured by Nanodrop. RNA integrity was verified on 1% agarose gel and by RNA Screen Tape assay (Agilent). cDNA was produced with High Capacity cDNA Reverse Transcriptase Kit (Applied Biosystems) or with qScript cDNA synthesis Kit (Quanta Bioscience) following the manufacturer’s instructions. cDNA was quantified on a real-time PCR detection system (CFX Connect; Bio-Rad) using iTaq universal SYBR Green Supermix (Bio-Rad). Cycling conditions consisted of initial denaturation (3 min) at 95°C, followed by 40 cycles of 10 s denaturation at 95°C and annealing/extension for 30 s at 60°C. Primer sequences are available in Supplementary Table 1. qPCR reactions were initiated in a final volume of 10 μL, containing 300 nM each of forward and reverse primers, and 25 ng of cDNA. Transcription levels were normalized to beta-actin and elongation factor 2 (eEF2) as internal controls. Experiments were repeated three times and are presented as fold change relative to the control. Calculations were performed using CFX Manager software (Bio-Rad).

### Library preparation for RNA-seq

Messenger RNA (mRNA) was enriched from 1 μg of total RNA by Poly(A) mRNA Magnetic Isolation Module (New England Biolabs) according to the manufacturer’s instructions. cDNA libraries were constructed using the NEBNext Ultra RNA Library Prep Kit (New England Biolabs) following the manufacturer’s protocol. Library concentration was measured by DNA High Sensitivity Kit (Invitrogen) on a Qubit fluorometer (Invitrogen). The High Sensitivity D1000 ScreenTape assay combined with the 2200 TapeStation system (Agilent) was used to assess the quality of the libraries.

### Silencing

For gene silencing by siRNA, 3134 cells were cultured for 24 h in Opti-MEM I Reduced Serum Media (Gibco) and transfected with 50 nM siRNAs using Lipofectamine 3000 (Invitrogen) according to the manufacturer’s instructions. siRNA sequences are listed in Supplementary Table 1. DNA transfection was performed using jetPRIME (Polyplus, France). Following transfection, cells were cultured for 24 h in DMEM supplemented with 10% FBS, 2 mM L-glutamine, 0.5 mg/mL penicillin-streptomycin and 1 mM sodium pyruvate, and for an additional 24 h in CSS-based media. After 48 h following transfection, cells were treated with hormone (100 nM Dex or EtOH) for 1 h, and RNA was collected.

### Klf4 chromatin immunoprecipitation (ChIP) assay and qPCR

3134 cells were cultured in CSS for 24 h and treated with either vehicle (EtOH) or 100 nM Dex for 2 h. Cells were cross-linked for 10 min at 37°C in 1% formaldehyde (Sigma) followed by quenching with 125 mM glycine for 10 min. Crosslinked cells were resuspended in RIPA buffer (10 mM Tris pH 8, 1 mM EDTA pH 8, 140 mM NaCl, 0.2% SDS, 0.1% DOC) supplemented with protease inhibitors, and sonicated for 38 cycles of 30 s ON and 30 s OFF (Bioruptor sonicator, Diagenode). Cleared chromatin was incubated overnight with 10 μg α-KLF4 (sc-20691). Complexes were washed twice with RIPA buffer, twice with high-salt buffer (10 mM Tris pH 8, 1 mM EDTA pH 8, 500 mM NaCl, 1% Triton, 0.2% SDS, 0.1% DOC), twice with LiCl wash buffer (10 mM Tris pH 8, 1 mM EDTA pH 8, 0.25 M LiCl, 0.5% NP-40, 0.5% DOC), and once in TE buffer (10 mM Tris, 1 mM EDTA, pH8). Crosslinks were reversed with 0.2 mg/mL Proteinase K overnight at 65°C. Purified DNA served as a template for qPCR. Primers used for qPCR amplification are listed in Supplementary Table 2. Fold enrichment was calculated using CFX Manager software (normalizing for the input and relative to negative control primers).

### 4C-seq

4C was performed as described (29, 30). Proximity ligation junctions, reflecting *in vivo* spatial proximity, were generated with DpnII (New England Biolabs), followed by circularization with Csp6I (Thermo Scientific). Chromosomal contacts with the TSS containing fragment were amplified with inverse PCR primers (Supplementary methods) and sequenced on Illumina 2000 platform.

### Computational analysis

#### 4C seq analysis

4C sequenced reads were sorted into different FASTQ files for each viewpoint and condition according to the bait sequences using a custom made Perl script. Data were analyzed using the 4Cseqpipe program (31). Briefly, the algorithm calculates medians of normalized coverage for running windows of linearly increasing size (2–49 kb, 1 kb steps), within an X kb window surrounding the bait. For each (2-49) window, the procedure generated a normalized signal for each 1 kb across X kb, giving rise to a table with 48 columns and X rows. Cell i,j in the table represents the normalized signal in the i-th range (2-49 kb) in the j-th position on the chromosome (from the X kb surrounding the bait, each position is 1 kb). These windows are presented in the figures as color-coded multiscale diagrams. In addition, a high resolution contact intensity trend line, depicting the medians of 5 kb running windows, is presented together with the 20th and 80th percentiles. To define contact domains, we first assigned to each 1 kb j position the maximum i-th range (2-49 kb) with a normalized signal higher than the top quartile of all normalized signals. Contact domains were obtained by merging j positions that were assigned with the top i-th range (49 kb), meaning their normalized signal is higher across all ranges, and closer than 15 kb.

#### Analysis of ChIP-seq and DNase I-seq data

GR, AP-1, RNA Pol-II, EP300 ChIP-seq and DNase I-seq data from 3134 cells treated with Dex for 1 h from (1, 2, 32) (SRP004871, SRP007111, GSE61236) was analyzed as follows. Sequenced reads were aligned to the mouse genome (mm9) using BOWTIE (33), and peaks were called using MACS2 (34) with default parameters for ChIP, and –nomodel; --shift -100; --extsize 200 parameters for DHS. A p-value cutoff of 1e-10 was used for AP1, CTCF and P300; 1e-4 for GR; and 1e-3 for DHS.

#### Pol-II ChIP-seq analysis

A Pol-II density signal was assigned for each gene. Genes were considered to have detectable expression if the normalized density signal was greater than 0.5 (median expression) under either of the conditions (10,565 expressed genes). The ratio (log_2_ fold change) between vehicle and Dex treated cells was calculated, and z-score normalization was applied to the log_2_FC values. Genes were scored as GR-regulated if the absolute value of Dex-dependent z-score was greater than 2 standard deviations.

#### RNA seq analysis

Reads were aligned to the mouse genome using STAR (35), and were counted on genes using HTseq (36). For each factor that was silenced four combination of enrichment analysis were done – si-Neg.C. with and without Dex, target genes siRNA with and without Dex, si-Neg.C. and gene-specific siRNA without Dex and si-Neg.C. and specific siRNA with Dex. Enrichment analysis between each combination was done using edgeR that assigns to each gene a FDR value and calculates the logFC between the two conditions (37). Genes with FDR <0.05 and logFC greater than |0.5| were considered differentially expressed.

#### Motif analysis

Motif discovery analysis was performed by the Homer suite, using the “findMotifsGenome.pl” program(38) with the paramters “-size given -N 20000 -cpg –noweight” for both DHS and GBS. Regions of local maxima of read count within DHS peaks (sub-peaks) were calculated using a custom R script. 100-250 bp regions with the highest read count in the DHS peaks, and 200 bp centered at the GR peak summits were submitted to Homer. Motif analysis was done for DHS or GR peaks within the domains of GR up regulated genes (up domains) or GR down regulated genes (down domains) with whole genome as background (HOMER default). The relative motif enrichment in up domains or down domains was calculated as the ratio between the proportions of the motifs in each group. The most highly enriched motifs are presented in Figures 5 and 6.

**Figure 1.**
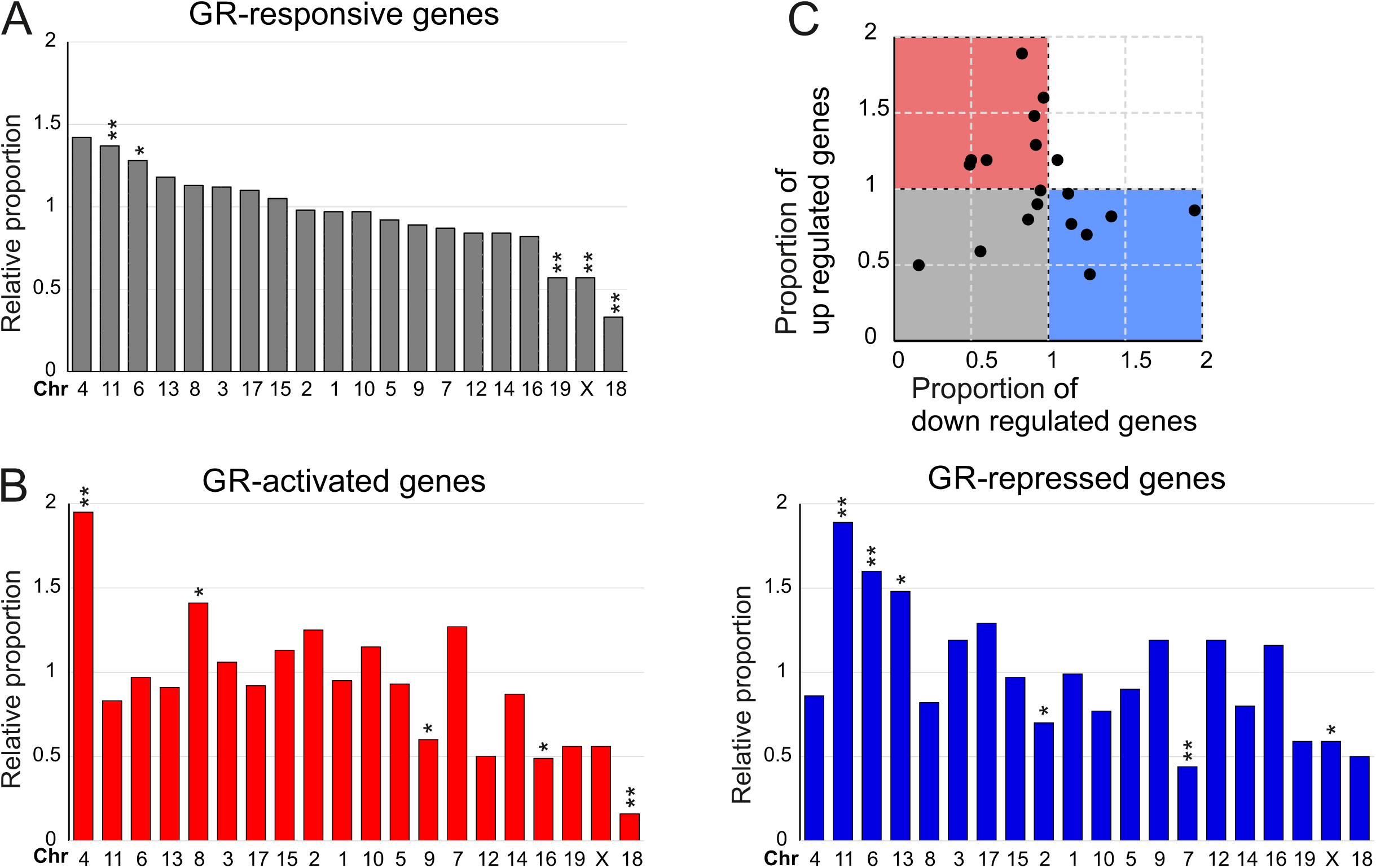
Non-random distribution of the transcriptional response to GR across the mouse genome. (**a**) Chromosome-specific enrichment of GR-responsive genes (proportion of GR-responsive genes on the chromosome /proportion of genes on the chromosome). Chromosomes were aligned according to their degree of enrichment. (**b**) Chromosome-specific enrichment for GR-activated (red) or GR-repressed (blue) genes. Chromosomes were aligned according to the order in A. **p<0.05, *p<0.1, proportional test. (**c**) The relative proportion of GR-upregulated genes plotted against the relative proportion of GR-down-regulated genes for each chromosome.

**Figure 2.**
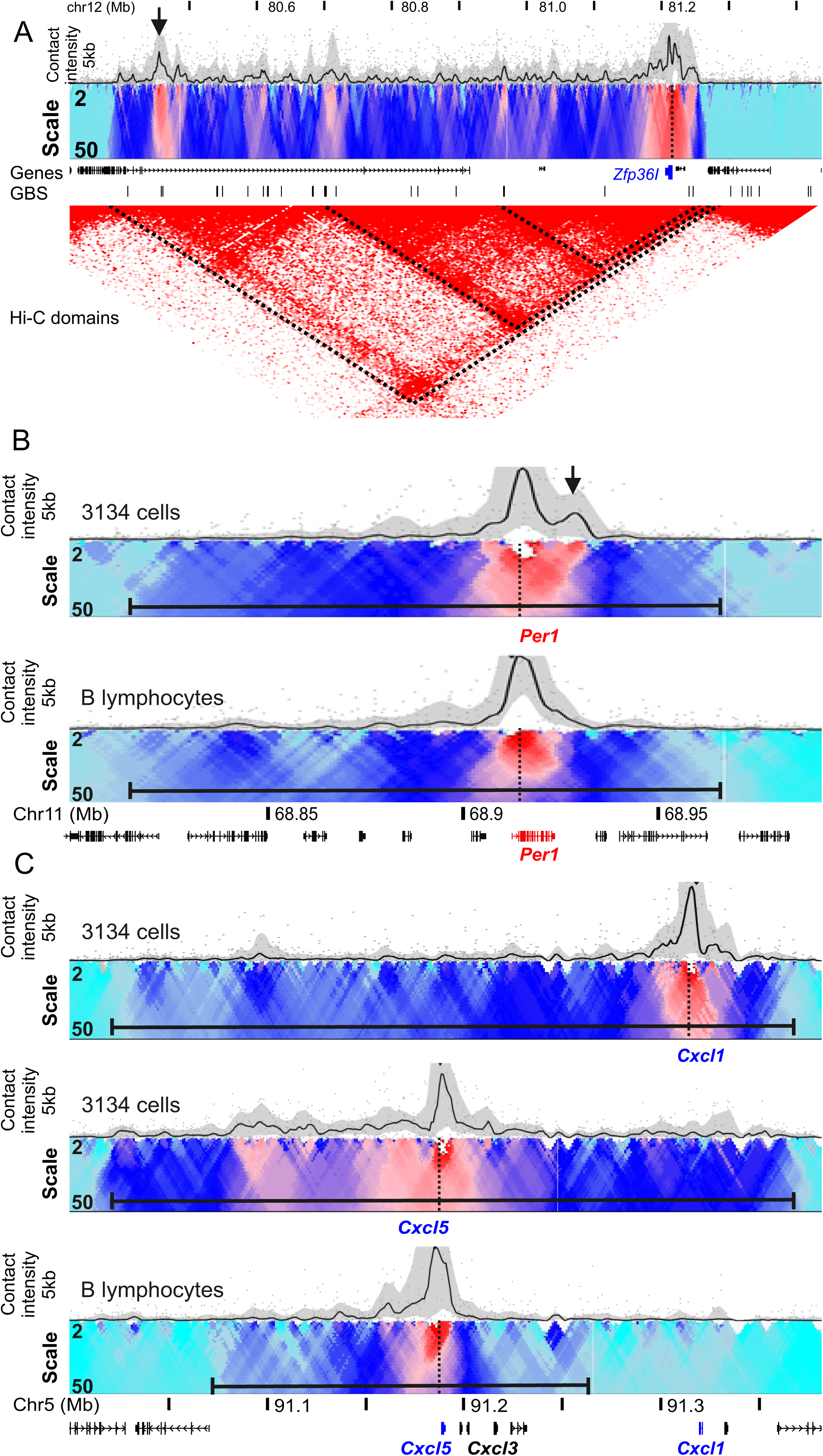
Spatial domains of GR-responsive genes. (**a**) High resolution chromosomal contact (4C-seq signal) profile of *Zfp36I* GR-repressed gene (depicted in blue) in 3134 cells. 4C contact profiles with the viewpoint (highlighted by a vertical dashed line) are shown in a trend line (upper panel, black line, using 5kb sliding window), and in a color-coded scale domainogram which shows relative interactions (red indicates the strongest interactions) in a sliding window ranging from 2 to 50 kb. Hi-C domains from murine CH12-LX cells 76 are shown. GBS –GR binding sites from ChIP. (**b**) 4C-seq profile of *Per1* GR-induced gene (depicted in red) in mammary 3134 cells and B lymphocytes. Arrow indicates cell type-specific contact (**c**) 4C-seq profile of *Cxcl5* and *Cxcl1* GR-repressed genes (depicted in blue) in mammary 3134 cells, and *Cxcl5* in B lymphocytes. The TSS fragment of the indicated genes was used as 4C bait. The horizontal black bar indicates the 4C spatial domain, and GR responsive genes are marked in blue. Genomic mm9 coordinates are given.

**Figure 3.**
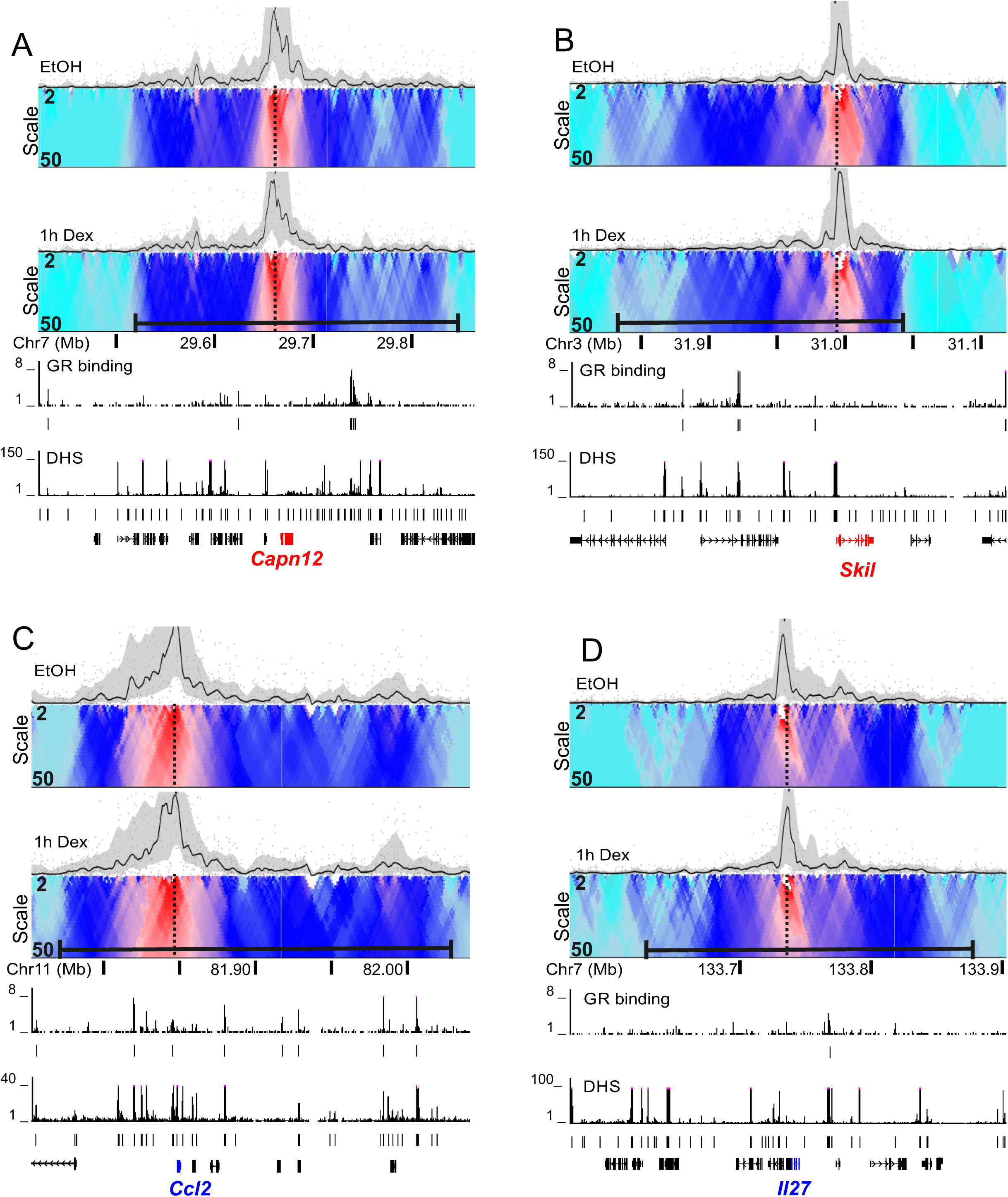
Stable chromosomal topology of GR-responsive genes. Spatial domains of GR-activated genes *Cpan12* (**a**) and *Skil* (**b**), as well as GR-repressed genes *Ccl2* (**c**) and *Il27* (**d**) are similar in vehicle (EtOH) and Dex (100nM, 1h) treated cells. GR-binding sites (ChIP-seq), and accessible regulatory sites (DNase-seq) from hormone treated cells (100nM, 1h) are indicated. GR-activated and repressed genes are marked in red and blue, respectively. Genomic mm9 coordinates.

**Figure 4.**
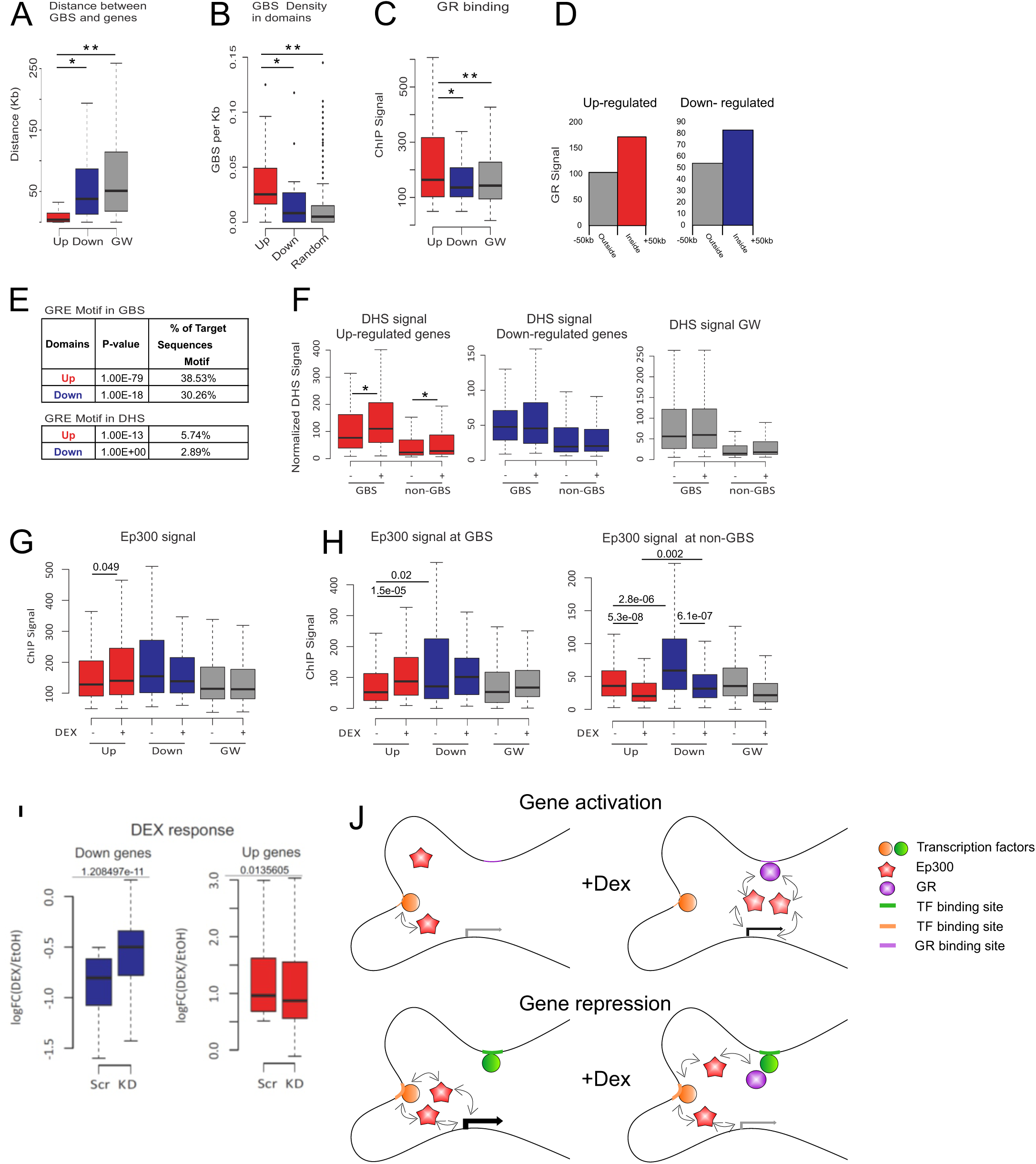
Features of regulatory elements associated with GR-regulated genes. (**a**) Boxplot showing the distance between TSS and the nearest GBS of GR up-regulated (red), down-regulated (blue), and genome-wide (GW) transcribed genes (grey). *p=3.2e-34, ** p=2.9e-76, Wilcoxon test. (**b**) GBS density (peaks per kb) in 4C domains of GR up-regulated (red), down-regulated (blue) and GW (random 200 kb domains, grey). *p=0.006, **p=1e-9, Wilcoxon test. (**c**) GR ChIP signal at GBS associated with up-, down-regulated genes and genome-wide (grey). *p=5.7e-6, **p=8.6e-19, Wilcoxon test. (d)Profiles of aligned 4C domain border regions (left and right borders combined) are shown for GR binding (sum of ChIP signal in 50kb from border). Red/blue bar and positive coordinates, inside 4C domains; gray bar and negative coordinates, outside 4C domains. (**e**) GRE motif in GBS and DHS associated with promoters of up- and down-regulated genes. (**f**) Chromatin accessibility (DHS) signal in GBS and non-GBS regulatory sites within domains of GR up-regulated (red), down-regulated (blue) and genome (grey), before (-) and after (+) 1h Dex treatment. *p<0.005, Wilcoxon test. (**g**) Ep300 signal at peaks within 4C domains. *p=0.049, Wilcoxon test. (**h**) Similar analysis for Ep300 loci at GBS and accessible sites where GR does not bind (non-GBS). p Wilcoxon test is indicated. (**i**) siRNA transfected cells treated with vehicle (- Dex) or Dex. p-values are indicated, Wilcoxon test. (**j**) A model for gene regulation by cross talk between regulatory elements within defined spatial domain. In gene activation, direct GR binding to DNA increased Ep300 signal at GBS, Ep300 is also sequestered from weak enhancers (orange). In gene repression, GR is tethered by a transcription factor such as AP1 (green) and sequesters Ep300 from strong active enhancers (orange) within the defined spatial domain.

**Figure 5.**
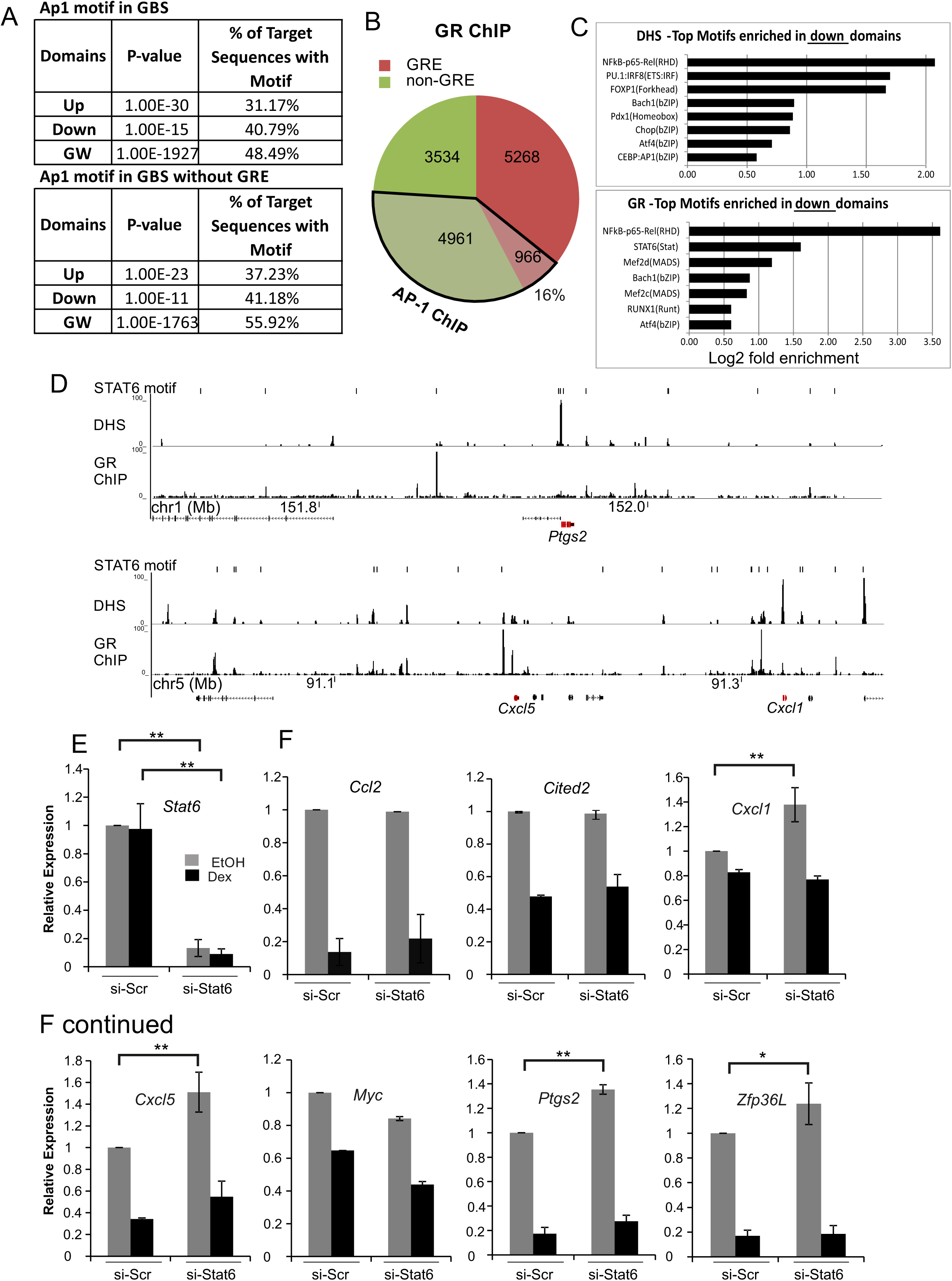
Transcription factor configuration at regulatory elements associated with GR-repressed genes. (**a**) AP-1 motif in GBS (with or without GRE) associated with GR up- or down-regulated genes and genome-wide (GW). (**b**) GR binding loci from ChIP-seq with (red) or without (green) GRE sequence are shown. AP-1 binding loci from ChIP-seq that overlap GBS are indicated in black frame. (**c**) Motifs in DHSs and GBSs associated with GR repressed genes, enriched relatively to activated genes. (**d**) Recognition motifs of STAT6 at DHS and GBS (ChIP-seq) (data from hormone (100nM Dex, 1h) treated cells) within 4C domains of *Ptgs2* and *Cxcl5* (depicted in red). Genomic mm9 coordinates. (**e**) Down regulation of STAT6 by siRNA. (**f**) Transcriptional response to GR activation in 3134 cells transfected with scrambled siRNA (si-Scr) or STAT6 siRNA. Error bars indicate SD of three biological repeats. * p<0.05, **p<0.01 t test.

**Figure 6.**
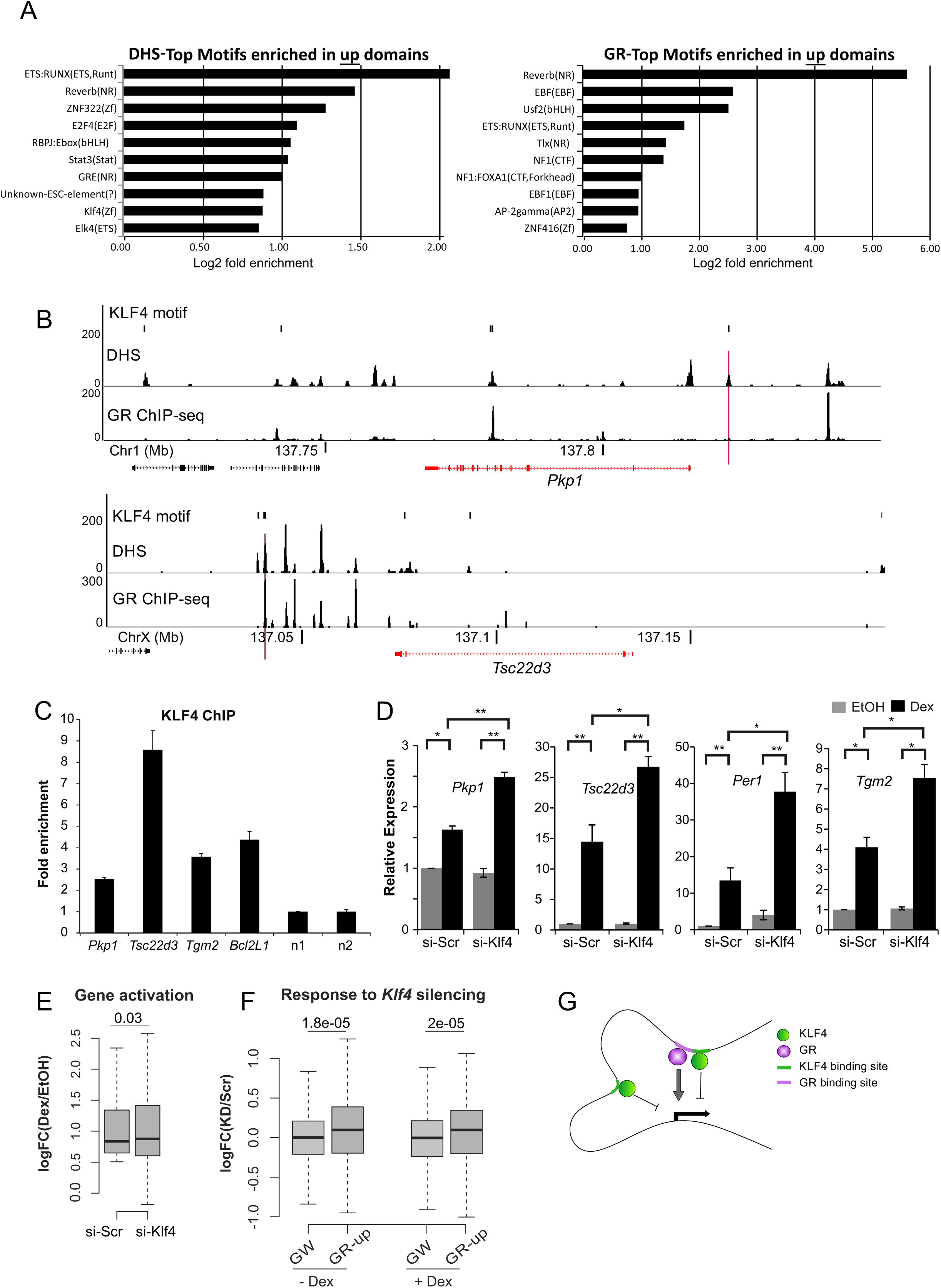
Transcription factor configuration at regulatory elements associated with GR-activated genes revealed Klf4. (**a**) Motifs in DHSs and GBSs associated with GR activated genes, enriched relatively to repressed genes. (**b**) Recognition motifs of Klf4 at DHS and GBS (ChIP-seq) (data from hormone treated cells (100nM Dex, 1h)), within 4C domains of *Pkp1* and *Tsc22d3* (depicted in red). Genomic mm9 coordinates. vertical red lines indicate loci that were validated by Klf4 ChIP. (**c**) KLF4 binding by ChIP-qPCR to regulatory elements with the Klf4 motif (indicated by vertical red line in Figure 6B, S3) n=negative control loci. (**d**) Transcriptional response to GR activation (1h Dex) measured by RT-qPCR in 3134 cells transfected with scrambled (si-Scr) or Klf4 siRNA. Error bars indicate SD of three biological repeats. * p<0.05, **p<0.01, Student’s t test. (**e**) log2 of the fold change of genes activated by Dex in 3134 cells transfected with scrambled (si-Scr) or Klf4 siRNA. (**f**) Transcriptional changes (log2 fold change Klf4 siRNA/ scrambled siRNA) of GR upregulated genes (GR-up) in cells treated with vehicle (-Dex) or Dex. The genome-wide (GW) changes are shown as control. p-values are indicated, Wilcoxon test. (**g**) A model for gene regulation by GR and KLF4. KLF4 binding to GRE and non-GRE elements within a defined spatial domain reduce gene activation by GR.

#### Enrichment and statistical analysis

The relative enrichment of GR-responsive, up- and down-regulated genes on specific chromosomes was calculated as the ratio between the fraction of GR-regulated genes (from all GR regulated genes) on the specific chromosome and the percentage of genes (from the entire genome) on that chromosome. The significance of the relative enrichment/depletion of genes was calculated using upper/lower tail proportional test using ‘prop.test’ functions in R.

The closest GBS to each TSS and its distance was calculated using Bedtools ‘closestBed’ script (39). Overlap between ChIP peaks and/or DHS and/or domains was identified using Bedtools ‘intersectbed’ script. Normalized ChIP signal was calculated as reads per million reads (from the whole dataset) per kb. Calculation of the overall GR ChIP signal in the 4C domains and flanking regions was performed by summing the ChIP signal in the 50 kb flanking both ends of all the 4C domains. Mann-Whitney-Wilcoxon Test statistics were applied to compare between the differences of each group in up or down domains versus the entire genome using R ‘wilcox.test’ function. The overlap between the GBS and GRE motifs (Homer motif bed file with all putative GREs in the genome) or nGREs (CTCCGGAGA, CTCCNGGAGA and CTCCNNGGAGA) was calculated using Bedtools ‘intersectbed’ script.

## RESULTS

### Spatial clustering of GR-responsive genes on mouse chromosomes

Hormone-activated GR translocates to the nucleus, where it binds to chromatin target sites and directly reprograms the transcription of multiple genes. To study the direct effects of GR on gene transcription, we screened for variation of activated (S5 phosphorylated) RNA polymerase-II occupancy one hour after hormone induction, using available ChIP-seq profiles in 3134 cells (32). We retrieved 263 up-regulated genes (>two-fold Pol-II occupancy along gene body regions, Z-score >2) and 251 down-regulated genes (<two-fold, Z-score <-2). The transcriptional response was validated by qRT-PCR (Supplementary Fig. 1)

Mammalian genomes are organized in hierarchies of chromosomal environments that can be defined by the frequency of specific long-range chromosomal associations. We previously showed that inter-chromosomal compartments, which represent higher level of cell-type specific chromosomal organization, are not particularly enriched for GR-regulated genes, or for a specific transcriptional response to GR (i.e. gene activation or repression) (40). Since each chromosome occupies a sub-nuclear volume termed chromosome territory (CT) (41), intra-chromosomal interactions are more frequent than inter-chromosomal ones. Interestingly, 6 out of 20 mouse chromosomes were either enriched (Chr 4, 6 and 11) or depleted (Chr 18, 19 and X) of GR-responsive genes (Fig. 1a). Notably, the enrichment was higher (11 out of 20 chromosomes) for a specific transcriptional response of activation or repression (Fig. 1b). The bias towards activation or repression in individual chromosomes is further notable as none of the mouse chromosomes is enriched for both responses (Fig 1c, white square). This non-random distribution of GR-responsive genes may facilitate their spatial clustering at the level of CT.

### Clustering of GR-responsive genes within spatial domains

Next, we sought to characterize in detail the spatial organization of GR-responsive genes within chromosomes using 4C-seq technology that measures chromosomal associations with a point of interest (29, 31). To understand how GR-responsive genes are regulated, we applied 4C-seq to define long-range associations of their transcription start sites (TSS). Interestingly, 4C-seq revealed that GR-responsive genes are embedded in domains of high spatial connectivity, which drop sharply at the edges (Fig. 2a). The 4C domains are in agreement with the partitioning of the chromosome to topologically associating domains (TADs) measured by Hi-C and 5C (42, 43) and nested neighborhoods or loops captured by high resolution Hi-C and ChIA-PET (44, 45). For example, the promoter of *Zfp36I* loops across two Hi-C sub-domains to a GBS located ∼700 kb downstream (Fig. 2a, arrow). Similarly to topological domains, 4C domains in mammary 3134 cells may exhibit conservation in other cell types (*Per1*, Fig. 2b). However, we also noted differences in the 4C domains between mammary 3134 cell line and murine B cells (*Cxcl5* locus, Fig. 2c). This could reflect variability in intra-domain contacts or in the insulation of the domain from its flanking regions (46). Importantly, the high resolution 4C revealed cell type-specific long range contacts within the domains of GR-regulated genes (Fig. 2b, arrow). For high genomic coverage, we measured (by 4C) the chromosomal contacts of 25 GR-activated and 15 GR-repressed genes. These genes have a wide range of transcription levels, and are spread across 13 mouse chromosomes (Supplementary Table 2). Interestingly, the domains that were captured by GR-activated genes include 12 additional GR activated genes (out of total 167 genes in the domains; p<10E-6, multinomial distribution). Moreover, three additional repressed genes are positioned within the domains of GR repressed genes (out of 57 genes in total; p<0.01, multinomial distribution). To validate the clustering of GR-responsive genes in spatial domains, we analyzed the chromosomal interactome of the GR-repressed gene *Cxcl3*, which is located in the domain of *Cxcl5*. Indeed, both genes share the same domain and similar contacts within the domain (Fig. 2c). We conclude that GR-responsive genes are embedded within spatial domains that are specialized for transcriptional induction or repression by hormone-activated GR.

### Dynamics of chromosomal structure in response to GR activation

GR activation elicits complex dynamic transitions in chromatin accessibility and enhancer activity (2, 32, 47, 48). Our analysis suggests that this ensemble of local transitions at regulatory elements converges in 3D with gene promoters to reprogram transcription of glucocorticoid-responsive genes. We therefore studied the dynamics of such long-range chromosomal interactions between TSS of GR-target genes and their entire scope of regulatory elements. We performed 4C for 11 GR-activated and 7 GR-repressed genes in vehicle (EtOH) or 1 h hormone (Dex) treated cells. Not surprisingly, the borders of the domains remained stable after Dex treatment. Moreover, despite robust transcriptional reprogramming by GR, chromosomal contacts within domains also remained stable, overall. These contacts reflect possible associations between promoters and enhancers, as depicted by DNase I hypersensitivity (DHS), GR and Ep300 binding (Fig. 3, Supplementary Fig. 2). These results are in line with our previous report of a stable chromosomal loop between the GR binding site at the *Lcn2* promoter and the distal *Ciz1* gene (24), and with global Hi-C study of the rapid (1 h) transcriptional response to TNF-α signaling (25).

### Building a functional catalogue of GBSs

GR-responsive genes cluster in 3D with multiple GR binding sites (GBS) and other non-GBS regulatory elements identified by their accessibility to DNase I digestion (Fig. 3). We hypothesized that these DNase I hypersensitive sites (DHSs) which are occupied by various transcription factors (TFs), may collaborate with GR to regulate gene expression. Importantly, the fact that the transcriptional response within a given spatial domain is specific to either activation or repression by GR, suggests that regulatory elements within the domain are specialized for either transcriptional response. This specialization suggests that these regulatory elements may serve as the toolbox to elicit specific transcriptional responses to GR signaling. Therefore, to uncover the mechanistic basis of transcriptional activation and repression by GR, we searched for features that discriminate between GBSs and DHSs of GR-activated and repressed genes. Using the 4C domains, we pooled catalogs of 231 GBS and 910 DHS sites associated with GR-activated genes, and 76 GBSs and 328 DHSs within the spatial domains of GR-repressed genes.

### Distinct configuration of chromatin features in GR-activated versus repressed domains

Although TF binding sites are, in general, distant from genes, we noted a marked difference in their distribution within domains of up- and down-regulated genes. As reported in several cell types (21, 49, 50), GBSs are closer to promoters of GR-activated genes compared to repressed genes (Median distance of 4 kb, 38 kb, and 51 kb from activated-, repressed- and genome-wide expressed genes, respectively. Figure 4a). Moreover, the defined 4C domains (see materials and methods) allowed us to go beyond the nearest GBS and consider all GBSs within the chromatin domain of the regulated gene. This revealed that both GR binding site density and *in vivo* ChIP signal are higher in domains of GR-activated relative to GR-repressed genes (Fig. 4b, c). Nevertheless, the GR signal is higher within domains of both classes compared to their flanking genomic regions (Fig.4d), indicating that the enrichment of GR binding within domains of GR-responsive genes is specific. The difference in ChIP signal corresponds to the higher proportion (38% in up domains and 30% in down domains) of GR binding sites with a canonical GR recognition motif (GRE) in domains of activated genes (Fig. 4e), suggesting that indirect GR binding is more pronounced at GBSs associated with GR-repressed genes.

Regulatory elements of the two groups also differ in chromatin structure and dynamics. Chromatin accessibility in GR-activated genes was increased following Dex treatment for both, GR binding sites as well as other, non-GR regulatory elements (Fig. 4f). Conversely, chromatin accessibility in GR-repressed genes was persistent after Dex treatment, both for GR and non-GR regulatory elements (Fig. 4f). Previous studies have shown that GR sites are overall accessible prior to GR activation and that this accessibility increases in a subset of loci following GR binding (2, 5, 10, 32, 51–53). Our results here show that hormone treatment elevates chromatin accessibility specifically at GR sites associated with GR-activated genes.

Next, we sought to determine whether the bi-polar transcriptional responses to GR are related to variations in enhancer activity by analyzing the loading of Ep300, which marks active enhancers (5, 32). Analyzing the Ep300 ChIP-seq signal at all the regulatory sites revealed increased signal specifically at enhancers associated with GR-activated genes (Fig. 4g). As expected, this increase can be attributed to increased *in vivo* binding of GR to its binding sites (p<1.2e-6, Fig. 4h). However unexpectedly, GR binding did not increase the Ep300 enhancer signal at sites associated with repressed genes (p<0.4, Fig. 4h). Importantly, although in both types of domains the Ep300 signal at non-GR sites was reduced following Dex treatment, the net reduction in the repressed domains was greater (from median of 59 to 31) than in the activated domains (from median of 36 to 20).

Altogether, GR binding sites associated with repressed genes are characterized by a low proportion of direct GR binding and higher distance from target genes relative to GR sites near activated genes. Importantly, GR binding at these sites does not increase the Ep300 signal, but leads to a decrease in the loading of this co-activator binding to non-GR regulatory elements, suggesting that gene repression by GR is predominantly passive, and is achieved through sequestering of Ep300 from other, non-GR binding sites and enhancers (Fig. 4g).

### Transcription factor composition of regulatory elements associated with GR-repressed genes

The distinct features of regulatory sites associated in 3D with GR-activated versus GR-repressed genes suggest that the molecular makeup of these two cistromes is different. The low frequency of GRE and low GR ChIP signal at GR sites which loop over promoters of repressed genes, suggest that GR binding at these sites is indirect. By interacting with other transcription factors, GR can bind to genomic regions bearing only the recognition motif of the partnering TF. However evidence exists for the direct repression by GR binding to negative GREs (nGRE) and recruitment of co-repressors such as SMRT/NCoR (15, 16). In agreement with previous studies (14, 18), we found that the nGRE motif is not sufficient to discriminate between activating and repressing GBSs since it is present at similar frequencies in both groups of GR sites (associated with up-regulated 0.12%, down-regulated 0.08%, and genome wide 0.08%).

Given that GR can repress genes by recruiting HDACs to GREs multiplexed with other transcription factors, such as IRF3 in macrophages (14), we searched for binding motifs of TFs that are enriched in GBSs within the spatial domains of repressed genes. Not surprisingly, this analysis revealed the activator protein 1 (AP-1) motif. However, the proportion of GBSs bearing an AP-1 motif was similar in down- and up-regulated domains (41% and 31%, respectively. Fig. 5a), indicating that AP-1 is not likely to play a differential role in gene activation and repression by GR. Although studies of individual target genes showed that GR represses AP-1-regulated genes by tethering to AP-1 and inhibiting its activity through a trans-repression mechanism (11, 54), genome-wide analysis of AP-1 binding in 3134 cells revealed that its activity is not restricted to GR-suppressed genes (1). According to Biddie et al., AP-1 has a global role in priming and maintaining accessible chromatin at GBSs, even prior to GR activation (1). Moreover, a similar proportion of GR-induced and -repressed genes are compromised by inhibiting AP-1 binding, suggesting that AP-1 is equally important for both transcriptional responses (1). Interestingly, combining GR ChIP-seq with AP-1 ChIP-seq revealed that the proportion of direct GR binding relative to the common binding sites is very low, as only 16% of genomic GBSs that overlap with AP-1 binding harbor a GRE (Fig. 5b). This proportion is maintained in the domains of GR repressed genes (16%, 5/32) while doubled, but still low, in the activated domains (30%, 27/88). Though one may assume that GR is tethered by AP-1 to the remaining loci, we found that only ∼40% of the genomic GRE-less GBSs, which are bound by AP-1, contain an AP-1 motif. This suggests that at the majority of common binding sites, both GR and AP-1 are tethered by additional factors.

To identify possible transcription factors that define regulatory elements associated with GR-repressed genes, we first performed motif discovery analysis in GR sites and DHSs using the genome as background. Then, for motifs with p value lower than 0.01, we calculated the ratio of motif in the “repressed” relative to the “activated” domains. This analysis motif enrichment specifically in the GR-repressing relative to GR-activating regulatory sites revealed nuclear factor kappa-B (NF-κB) and signal transducer and activator of transcription 6 (STAT6) motifs (Figure 5C). Notably, the AP-1 motif was not found to be enriched specifically in repressed domains, in line with the similar role of AP-1 in gene activation and repression by GR (1).

The enrichment of the NF-κB motif in the GBSs of the repressed genes is in line with many studies implicating GR-mediated repression of NF-κB target genes by a tethering mechanism (10– 13). We chose to study STAT6 as a potential novel global GR co-repressor. Previous studies have shown that GR physically interacts with several members of the STAT family in both gene activation and repression (9, 11, 47, 55). STAT6 interacts with GR in human CTLL2 T cells (56), and is involved in gene activation and repression by GR. Reporter constructs bearing a STAT6 motif were used to show that STAT6 and GR synergize to activate the β-casein promoter (57), while they mutually antagonize the expression of MMTV-LTR (56) in CTLL2 cells. In addition, GR was unable to inhibit transcription of three different STAT6-driven reporter constructs in airway epithelial cells (58), though it is possible that the mutual effect of GR and STAT6 is gene- and cell type-specific. In addition, reporter constructs may be restricted in their ability to reflect the regulatory complexity by GR, since they include a limited number of regulatory elements. To study the interplay between STAT6 and GR, we downregulated *STAT6* expression by siRNA, and measured the response of GR-repressed genes bearing the STAT6 motif in their associated GBSs (Fig. 5d, Supplementary Fig. 3). Downregulation of *STAT6* (Fig. 5e) led to elevated expression of *Cxcl5, Cxcl1, Ptgs2* and *Zfp36l1* prior to hormone application, suggesting that these genes are regulated by STAT6. However, these genes were negatively regulated by GR regardless of STAT6 inhibition (Fig. 5f). Thus GR and STAT6 may act independently as negative regulators of these genes.

### Deciphering the regulatory partners of GR in gene activation

To uncover transcription factors that are associated specifically with gene activation by GR, we calculated the ratio between the proportion of motifs in activated domains relative to those in repressed domains. Analysis of both GBSs and DHS, uncovered motifs of candidate factors that are linked to GR biology, but for which less is known about their co-regulatory activity at the chromatin level. Notably, analysis of DHSs associated with GR-induced promoters revealed factors that may be overlooked by analyzing GR sites alone.

### Klf4 counteracts gene activation by GR

Our focused motif discovery revealed the Krüppel-like factor 4 (Klf4) binding sequence in DHSs preferentially associated with promoters of GR-induced genes relative to DHSs of the GR-repressed loci (Fig. 6a,b, Supplementary Fig. 4a). The computational prediction of Klf4 binding was confirmed by Klf4 ChIP-qPCR for GBSs and non-GBSs in the domains of *Tsc22d3, Tgm2, Bcl2l1*, and *Pkp1* genes (Fig. 6b,c, Supplementary Fig. 4a). This suggests that Klf4 may regulate GR-target genes by binding to GBSs, and/or by associating with other, non-GBS regulatory elements. To analyze the contribution of Klf4 to the transcriptional regulation by GR, we silenced Klf4 by siRNA and treated the cells with Dex. Transcriptional activation of *Tsc22d3, Tgm2, Bcl2l1*, and *Pkp1* genes was enhanced in cells expressing Klf4 siRNA relative to cells expressing control siRNA (Fig. 6d, Supplementary Fig. 4b). Moreover, RNA-seq analysis showed global enhancement of the transcriptional activation by GR under Klf4 silencing (Fig. 6e, Supplementary Fig. 4c,d,e). This suggests that indeed, Klf4 co-regulates the transcriptional response of these genes to GR by antagonizing their GC-mediated transcriptional activation (Fig. 6g). Notably, in line with the repressive effect of Klf4 on GR-target genes, their transcription was also elevated by Klf4 silencing also prior to Dex application (Fig. 6f).

### Rev-erb and ROR

The Rev-erb motif, which was highly enriched in GBSs associated with activated genes (Fig. 6a), is also the recognition sequence of ROR (RAR related orphan receptors), termed ROR response element (RORE). Rev-erb and ROR belong to the nuclear receptor super family of transcription factors. The activities of the two factors are thought to be antagonistic to each other. Rev-erb represses transcription by recruiting the NCoR/SMRT co-repressor, while RORs recruit the SRC and/or EP300 co-activators to ROREs (59). Rev-erb and ROR factors are part of the circadian clock mechanism and are involved in regulating many physiological processes, including metabolism, development and inflammatory processes (59). Notably, these same physiological processes are also regulated by glucocorticoids (GCs). Moreover, GCs are released from the adrenal glands in a circadian manner, and GR reset the circadian clock in peripheral tissues (60). These links between Rev-erb /ROR and GR biology, together with the enrichment of RORE at GBSs, prompted us to investigate the crosstalk between these factors in transcriptional regulation by GCs.

From the members of the Rev-erb and ROR families, we focused on ROR β, ROR γ and Rev-erb β (Nr1f2, Nr1Ff3 andNr1d2, respectively) which were the abundant transcripts in 3134 cells (Supplementary Fig. 5a). Since the gene targets of ROR and Rev-erb in 3134 cells are not known, we first analyzed nuclear factor IL-3 regulated (*Nfil3*) gene, which is induced by Dex in 3134 cells and is associated with five RORE motifs in 3D (Fig. 7a). In addition, previous studies have shown that Rev-erbα binds the promoter, and regulates the expression of in mouse liver, suggesting that *Nfil3* is a direct target of Rev-erbα in these cells (61). In order to study the contribution of ROR and Rev-erb factors to gene regulation by GR, we downregulated their expression by siRNA and measured the transcriptional response to Dex application. Importantly, in line with their predicted activating role, knocking down *RORβ* and *RORγ* in 3134 cells reduced the transcriptional induction of *Nfil3* (Fig. 7b, Supplementary Fig. 5c,e), suggesting that RORγ collaborates with GR in the up-regulation of this gene. Importantly, extending this analysis to the whole genome revealed similar trends (Fig. 7c). Moreover, ROR knock down did not affect the basal expression of GR-activated genes, but only had an effect on gene expression following Dex treatment (Fig. 7c), indicating that ROR collaborates with GR in gene activation in a hormone-dependent manner.

**Figure 7.**
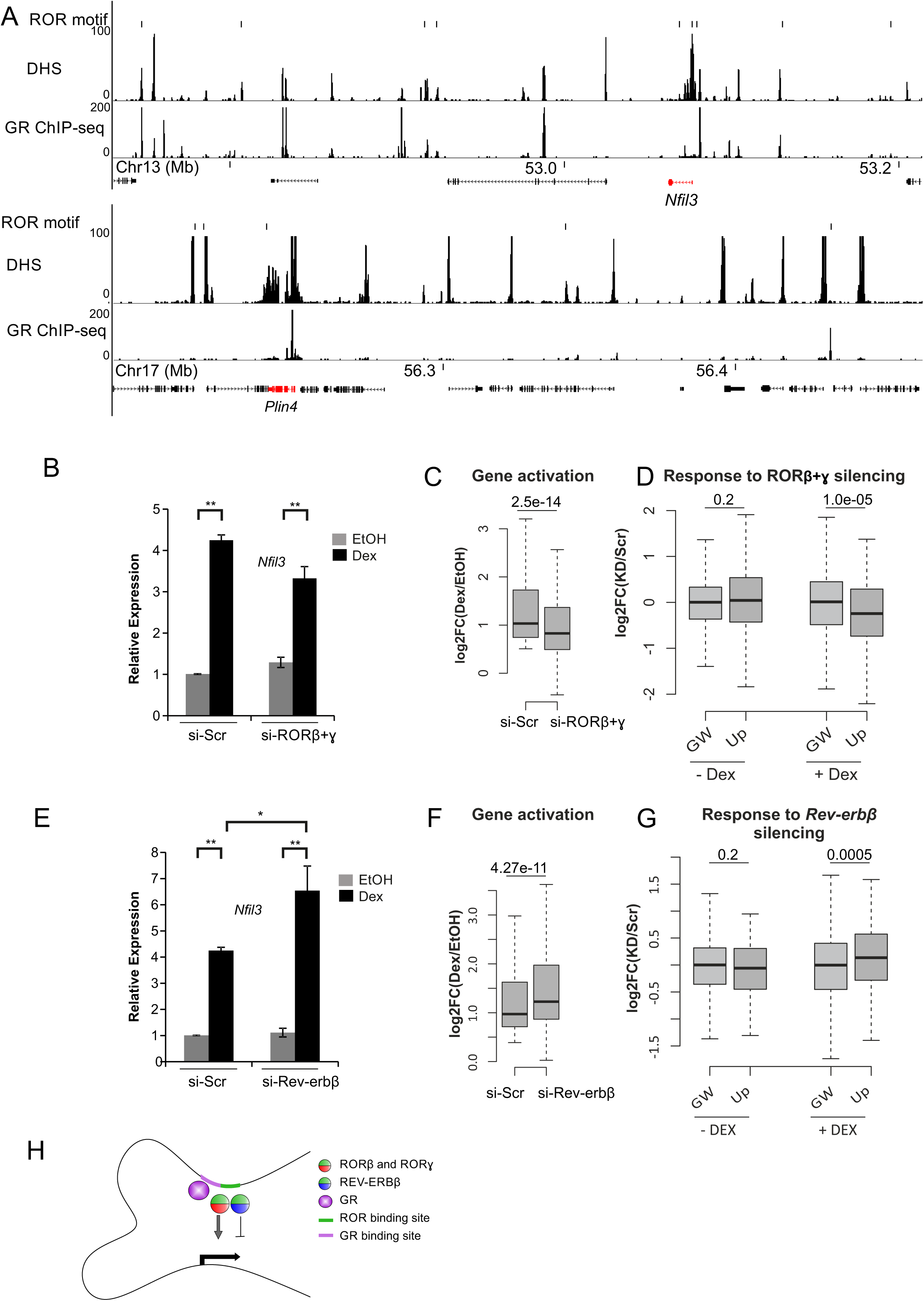
ROR, Rev-erb and GR cooperation. (**a**) Recognition motifs of ROR at DHS and GBS (ChIP-seq) (data from hormone treated cells (100nM Dex, 1h)) within 4C domains of *Nfil3* and *Plin4* (depicted in red). Genomic mm9 coordinates. (**b**,**e**) Transcriptional response to GR activation (1h Dex) in 3134 cells transfected with scrambled (si-Scr) or Rev-erbβ / RORβ and RORγ siRNA. Error bars indicate SD of three biological repeats. * p<0.05, **p<0.01 Student’s t test. (**c**,**f**) log2 of the fold change of genes activated by Dex in 3134 cells transfected with scrambled (si-Scr) or Rev-erbβ / RORβ and RORγ siRNA. (**d**,**g**) Transcriptional changes (log2 fold change knock down (KD)/scrambled (Scr) siRNA) of GR upregulated genes (GR-up) in Rev-erbβ (**d**) and RORβ and RORγ (**g**) siRNA transfected cells treated with vehicle (-Dex) or Dex. p-values are indicated, Wilcoxon test. (**h**) A model for gene regulation by GR, ROR and Rev-erb. Both, Rev-erbβ and RORβ/γ, negative and positive GR co-regulators, can bind a composite GR binding site harboring a GRE.

In line with its expected repressive activity, knocking down *Rev-erbβ* led to higher Dex-mediated activation of *Nfil3* (Fig. 7e, Supplementary Fig. 5b,d), indicating that Rev-erbβ antagonizes the rapid hormone-induced gene activation by GR. Genome-wide RNA-seq analysis shows that, indeed, transcriptional activation by GR is significantly increased under *Rev-erbβ* silencing (Fig. 7f) in a hormone-dependent manner (Fig. 7g).

Overall, our results suggest that Rev-erb and ROR are novel transcriptional co-regulators of GR (Fig. 7h).

## DISCUSSION

Despite our capacity to measure transcript levels with great accuracy, and to describe transcription factor binding loci and chromatin structure, it is not clear how transcription factors activate some genes, while at the same time repressing others. Hormone-activated GR provides an excellent model system, as opposed to multifactorial long-term responses, to address this question given its rapid, specific and direct transcriptional regulatory activity. Moreover, understanding the mechanistic basis of gene activation and repression by GR is important for appreciating its central immunosuppressive, metabolo-regulatory, and other physiological functions.

Indeed, the transcriptional response to GR is robust and rapid. 263 and 251 genes were up- or down-regulated, respectively, within one hour of Dex application. We noted that the genomic organization of GR responsive genes across the mouse genome is not random, to the extent that more than half of the mouse chromosomes are either enriched or depleted for a specific transcriptional response. Although previous studies identified non-random linear organization of co-expressed genes, clustering of co-regulated genes has been more challenging to characterize (62). Since each chromosome is confined to a given spatial territory, we propose that this chromosome-level enrichment of genes with similar transcriptional response to GR supports their spatial clustering, although these compartments are not exclusive to either response (40). However higher resolution study of the local organization of 40 GR-responsive genes by 4C-seq, revealed that they are embedded within structural domains, which are chromosomal segments characterized by high internal connectivity. Importantly, analysis of additional GR-responsive genes that were found to cluster within these domains revealed that the domains are exclusive to either activation or repression. Our findings suggest that these domains insulate the effect of the residing regulatory elements from other domains, and also include the required regulatory elements that bring about a specific transcriptional response. Like the structure of TAD, the 4C domains are overall similar in 3134 and B cells, and may contain nested loops (44–46).

The promoter-centric 4C-seq revealed that GR-responsive genes are engaged with multiple GBS, non-GBS regulatory elements, and other gene promoters from within the structural domain. These spatial clusters within the domains remain stable following GR binding and transcriptional activation or repression. Similarly, stable chromosomal associations were measured by 3C between the promoter of the GR-induced *Lcn2* gene and a 32 kb distant GBS in 3134 cells (24), and in mouse liver (63) as well, using a genome-wide Hi-C study that measured stable chromosomal associations between TNF-α responsive genes and enhancers following 1 h TNF-α treatment (25). On the other hand, there are examples of dynamic promoter-enhancer loops in response to prolonged activation of other signal-responsive factors. Chromosomal contacts measured by Hi-C were enhanced or reduced in concordance with the transcriptional response to progesterone receptor (PR) activation for 6 h in T47D cells (26), and also during early differentiation when measured by promoter capture Hi-C 4h following activation of multiple adipogenesis promoting transcription factors in 3T3-L1 cells (64). In another study, chromosomal contacts, measured by 4C-seq, were enhanced at a subset of EP300 binding loci following activation of GR and/or NF-κB in HeLa cells (8). Additional studies are required to determine whether dynamic loops, which were also reported in 3134 cells (32), are the exception or the rule, or whether their formation is related to the extent of activation, cell type, treatment or transcription factor. Careful inspection of dynamic loops from this and other reports suggest that they may represent enhancement or reduction of contact frequencies in pre-existing structures. This may require single-cell measurement by imaging to complement the information about relative changes from these cell population data.

The spatial partitioning of GR-induced and -repressed domains suggests that their regulatory elements dictate a specific transcriptional response to GR. This provided us a unique opportunity to uncover the differential molecular makeup of the bipolar (activation or repression) transcriptional responses to hormone-activated GR. Importantly, the ensemble of regulatory elements that converge in 3D include non-GBS regulatory loci that may collaborate with GBSs to define a specific transcriptional response.

Our analysis of GBSs and DHS associated with GR-repressed genes indicated that molecular sequestering is a major mechanism for GR-mediated repression. GBSs in the repressed domains are characterized by relatively low density, low frequency of the GRE motif, and low GR ChIP signal, suggesting that GR binding at these sites is indirect via tethering to an intermediate DNA-bound transcription factor (18, 65, 66). The rapid gene repression in these domains can be explained by an immediate decrease of the Ep300 signal from the promoter-associated non-GBS loci following GC binding. In agreement with the proposed factor sequestering model of nuclear receptor-mediated gene silencing, we suggest that GR competes with non-GBSs for a limited amount of CBP/Ep300 (67), thereby impairing their positive activity and repressing their target genes. Surprisingly, GR binding to chromatin also sequestered Ep300 from non-GRE regulatory elements associated with activated genes. However, since prior to Dex application, the Ep300 ChIP signal was higher at non-GRE enhancers of the repressed, relative to the activated genes, the magnitude of Ep300 depletion was greater in non-GRE enhancers of the repressed genes. In agreement, it was recently shown that high occupancy-enhancers are particularly sensitive to MED1 loss following TNFα treatment in adipocytes (68).

Moreover, we observed different dynamics of the Ep300 signal on activating and repressing GBSs. Following GR binding, the Ep300 signal remained stable at the GBSs of the repressed domains, while it dramatically increased on GBSs of GR-activated domains. We propose model by which this difference results from relatively greater dynamic binding of GR to chromatin in the repressed domains. GR interacts dynamically with other transcription factors, co-activators and the chromatin template (69, 70). In the repressed domains, the kinetics of any indirect GR binding to chromatin would be determined by its binding dynamics to the tethering factor (for example AP-1) and binding of the tethering factor to DNA. Consequently, although GR binding at the repressing GBSs does not alter the average Ep300 ChIP signal over the entire cell population, the net exchange rate at the repressing GBSs is rapid relative to direct binding of GR to DNA in the activated domains. Thus, even if the affinity between GR to Ep300 is similar in the activating and repressing GBSs, on average, a shorter residence time of GR on GBS by tethering is likely to be less productive than direct binding. In addition, the increased exchange rate would provide more chances for GR to sequester Ep300 from proximal non-GBS regulatory loci, allowing efficient sequestering using fewer GBSs in the repressed compartments. Importantly, this model implies that the local Ep300 concentrations within the spatial compartments of activated and repressed genes may be more significant than the global nucleoplasmic concentration for determining the available Ep300 for competition (Fig. 4i). It will be interesting to explore whether GR sequesters other co-activators such as SRC-1 and GRIP1 (7–10), and whether there are specific co-activator compositions on non-GBS enhancers associated with repressed genes.

The spatial and functional partitioning of GBSs and DHS provided an opportunity to uncover their differential transcription factor assignment. Surprisingly, regulatory elements in positive and negative domains are similarly decorated with AP-1 binding sites, suggesting that globally, despite its tethering and priming capacity, AP-1 does not dictate a specific transcriptional response. While several transcription factor motifs are enriched in negative domains, knock-down experiments for STAT6 suggest that these factors regulate genes within the domains independently of GR activity. This suggests that in a global manner, factor sequestering is a predominant mechanism for rapid gene repression by GR.

Since Ep300 concentration, availability and regulatory effect may be dictated by the number and ratios of GBSs to non-GBSs with different features within the local compartments of GR responsive genes, these parameters should be further investigated by perturbing different enhancers in their natural spatial context. Understanding the ensemble of features of regulatory elements and gene promoters in the context of the local compartment can provide a quantitative prediction for gene repression.

The GR-activated cistrome was enriched for motifs of several candidate GR cofactors. Klf4 is thought to cooperate with GR in mouse skin development since both factors regulate a similar set of genes (71). Klf4 binds to GBSs of *Tsc22d3* and *Zfp36*, but regulates only *Tsc22d3* expression in mouse keratinocytes (72). Here we show that Klf4 regulates multiple GR target genes in mouse mammary cells. Notably, the Klf4 motif is primarily enriched in non-GBS regulatory elements. This important result indicates that the complex gene regulation by GR includes spatial crosstalk between multiple GBS and non-GBS regulatory elements that cluster together within the three-dimensional space of the nucleus. Unexpectedly, Klf4 antagonizes GC-mediated transcriptional activation (Fig. 6g). Klf4 is a versatile transcription factor that possesses a transactivation and a repression domain. It is possible that gene co-activation or co-repression by Klf4 is regulated by post translational modifications in a cell-type and context-dependent manner (73).

Our results suggest that Rev-erb and ROR are novel co-regulators of GR. Their motif was enriched both in GBS and non-GBS regulatory loci associated with activated genes and knock-down experiments indicate that RORβ and RORγ collaborate with GR in gene activation, while Rev-erbβ showed co-repressor capacity (Fig. 7h). The antagonistic functions of Rev-erb and ROR may enable modular co-regulation of the rapid hormone-induced gene reprograming by GR. Rev-erb and ROR factors are widely expressed and the functions of each family member largely overlap, suggesting that Rev-erbs and RORs are core components of the rapid and dynamic gene regulation by GR in various cell-types. It is yet to be determined how the balance between Rev-erbs, RORs and glucocorticoid levels, which alternate in circadian oscillations, as well as being altered by stress, affect tissue- and gene-specific transcriptional levels. Moreover it was recently shown that ROR activity promotes chromatin decondensation, which facilitates subsequent REV-ERB loading in a “facilitated repression” model (74), suggesting that the interplay between ROR and REV-ERB is more than simple antagonism. Given that the majority of Rev-erbα binding sites in liver lack the RORE motif (75), it is possible that similarly to GR, the enrichment of RORE motif in the GR-activated genes may reflect differential mode of binding rather than differential binding to chromatin. Intriguingly it was recently shown that REV-ERBα binds GR together with HSP90 chaperone and thereby affects GR stability, nuclear localization and the expression of GR target genes (76). Thus it is tempting to speculate that the transcriptional activity of GR is also tuned by the interaction with REV-ERBβ in the cytoplasm of 3134 cells.

Finally, while the focus of this study was on gene activation and repression by GR, it is likely that the ensemble of regulatory elements within spatial domains, and their associated transcription factors integrate signals from several pathways and coordinate the magnitude of the transcriptional activity of genes within spatial domains across the mouse genome.

## ACCESSION NUMBERS

The data discussed in this publication have been deposited in NCBI’s Gene Expression Omnibus and are accessible through GEO Series accession number GSE106209. The data is in private status and the secure token to allow the assess of the data is unkvqssyrxyrzgh

## SUPPLEMENTARY DATA

Supplementary Data are available at NAR online.

## ACKNOWLEDGEMENT

A.S.P is supported by the Nehemia Levtzion Fellowship.

## FUNDING

This work is supported by Marie Curie Integration grant [FP7-PEOPLE-2013-CIG-618763 to O.H.] and I-CORE Program of the Planning and Budgeting Committee and The Israel Science Foundation [41/11 to O.H.]. O.H. and M.-H.S. are supported together by the United States-Israel Binational Science Foundation (BSF), Jerusalem, Israel [2013409]. This research is supported in part by the Intramural Research Program of the National Institutes of Health at the National Institute on Aging. Funding for open access charge: United States-Israel Binational Science Foundation (BSF).

## CONFLICT OF INTEREST

The authors have no competing financial interests.

## TABLE AND FIGURES LEGENDS

Figure 1. Non-random distribution of the transcriptional response to GR across the mouse genome. (a) Chromosome-specific enrichment of GR-responsive genes (proportion of GR-responsive genes on the chromosome/proportion of genes on the chromosome). Chromosomes were aligned according to their degree of enrichment. (b) Chromosome-specific enrichment for GR-activated (red) or GR-repressed (blue) genes. Chromosomes were aligned according to the order in A. **p<0.05, *p<0.1, proportional test. (c) The relative proportion of GR-upregulated genes plotted against the relative proportion of GR-down-regulated genes for each chromosome.

Figure 2. Spatial domains of GR-responsive genes. (a) High resolution chromosomal contact (4C-seq signal) profile of *Zfp36I* GR-repressed gene (depicted in blue) in 3134 cells. 4C contact profiles with the viewpoint (highlighted by a vertical dashed line) are shown in a trend line (upper panel, black line, using 5 kb sliding window), and in a color-coded scale domainogram which shows relative interactions (red indicates the strongest interactions) in a sliding window ranging from 2 to 50 kb. Hi-C domains from murine CH12-LX cells (76) are shown. GBS, GR binding sites from ChIP. (b) 4C-seq profile of *Per1*, GR-induced gene (depicted in red) in mammary 3134 cells and B lymphocytes. Arrow indicates cell type-specific contact (c) 4C-seq profile of *Cxcl5* and *Cxcl1* GR-repressed genes (depicted in blue) in mammary 3134 cells, and *Cxcl5* in B lymphocytes. The TSS fragment of the indicated genes was used as 4C bait. The horizontal black bar indicates the 4C spatial domain, and GR-responsive genes are marked in blue. Genomic mm9 coordinates.

Figure 3. Stable chromosomal topology of GR-responsive genes. Spatial domains of GR-activated genes *Cpan12* (a) and *Skil* (b), as well as GR-repressed genes *Ccl2* (c) and *Il27* (d), are similar in vehicle (EtOH) and Dex (100 nM, 1 h) treated cells. GR-binding sites (ChIP-seq), and accessible regulatory sites (DNase-seq) from hormone treated cells (100 nM, 1 h) are indicated. GR-activated and repressed genes are marked in red and blue, respectively. Genomic mm9 coordinates.

Figure 4. Features of regulatory elements associated with GR-regulated genes. (a) Boxplots showing the distance between TSS and the nearest GBS of GR up-regulated (red), down-regulated (blue), and genome-wide (GW) transcribed genes (white). *p=3.2e-34, ** p=2.9e-76, Wilcoxon test. (b) GBS density (peaks per kb) in 4C domains of GR up-regulated (red), down-regulated (blue) and GW (random 200 kb domains, white). *p=0.006, **p=1e-9, Wilcoxon test. (c) GR ChIP signal at GBS associated with up-, down-regulated genes and genome-wide (white). *p=5.7e-6, **p=8.6e-19, Wilcoxon test. (d) Profiles of aligned 4C domain border regions (left and right borders combined) are shown for GR binding (sum of ChIP signal in 50 kb from border). Red/blue bar and positive coordinates, inside 4C domains; white bar and negative coordinates, outside 4C domains. (e) GRE motif in GBS and DHS associated with promoters of up- and down-regulated genes. (f) Chromatin accessibility (DHS) signal in GBS and non-GBS regulatory sites within domains of GR up-regulated (red), down-regulated (blue) and genome (white), before (-) and after (+) 1 h Dex treatment. *p<0.005, Wilcoxon test. (g) Ep300 signal at peaks within 4C domains. *p=0.049, Wilcoxon test. (h) Similar analysis for Ep300 loci at GBS and accessible sites where GR does not bind (non-GBS). P value of Wilcoxon test is indicated. (i) siRNA transfected cells treated with vehicle (-Dex) or Dex. p-values are indicated, Wilcoxon test. (j) A model for gene regulation by cross talk between regulatory elements within defined spatial domain. In gene activation, direct GR binding to DNA increased Ep300 signal at GBS, Ep300 is also sequestered from weak enhancers (orange). In gene repression, GR is tethered by a transcription factor such as AP1 (green) and sequesters Ep300 from strong active enhancers (orange) within the defined spatial domain.

Figure 5. Transcription factor configuration at regulatory elements associated with GR-repressed genes. (a) AP-1 motif in GBS (with or without GRE) associated with GR up- or down-regulated genes and genome-wide (GW). (b) GR binding loci from ChIP-seq with (red) or without (green) GRE sequence are shown. AP-1 binding loci from ChIP-seq that overlap GBS are indicated in black frame. (c)Motifs in DHSs and GBSs associated with GR repressed genes, enriched relative to activated genes. (d) Recognition motifs of STAT6 at DHS and GBS (ChIP-seq) (data from hormone (100 nM Dex, 1 h) treated cells) within 4C domains of *Ptgs2* and *Cxcl5* (depicted in red). Genomic mm9 coordinates. (e) Downregulation of *STAT6* by siRNA. (f) Transcriptional response to GR activation in 3134 cells transfected with negative control siRNA (si-Neg.C.) or STAT6 siRNA. Error bars indicate SD of three biological repeats. * p<0.05, **p<0.01 t-test.

Figure 6. Transcription factor configuration at regulatory elements associated with GR-activated genes. (a) Motifs in DHSs and GBSs associated with GR activated genes, enriched relatively to repressed genes. (b) Recognition motifs of Klf4 at DHS and GBS (GR ChIP-seq) (data from hormone treated cells (100 nM Dex, 1 h)), within 4C domains of *Pkp1* and *Tsc22d3* (depicted in red). Genomic mm9 coordinates. Vertical red lines indicate loci that were validated by Klf4 ChIP. (c) Klf4 binding by ChIP-qPCR to regulatory elements with the Klf4 motif (indicated by vertical red line in Figure 6B, S3) n=negative control loci. (d) Transcriptional response to GR activation (1 h Dex) measured by RT-qPCR in 3134 cells transfected with negative control (si-Neg.C.) or Klf4 siRNA. Error bars indicate SD of three biological repeats. * p<0.05, **p<0.01, Student’s t-test. (e) Log2 of the fold change of genes activated by Dex in 3134 cells transfected with negative control (si-Neg.C.) or Klf4 siRNA. (f) Transcriptional changes (log2 fold change Klf4 siRNA/ negative control siRNA) of GR upregulated genes (GR-up) in cells treated with vehicle (-Dex) or Dex. The genome-wide (GW) changes are shown as control. P-values are indicated, Wilcoxon test. (g) A model for gene regulation by GR and Klf4. Klf4 binding to GRE and non-GRE elements within a defined spatial domain reduce gene activation by GR.

Figure 7. Cooperation between ROR, Rev-erb and GR transcription factors. (a) Recognition motifs of ROR and Rev-erb at DHS and GBS (ChIP-seq) (data from hormone-treated cells (100 nM Dex, 1 h)) within 4C domains of *Nfil3* and *Plin4* (depicted in red). Genomic mm9 coordinates. (b,e) Transcriptional response to GR activation (1 h Dex) in 3134 cells transfected with scrambled (si-Neg.C.) or Rev-erbβ / RORβ and RORγ siRNAs. Error bars indicate SD of three biological repeats. * p<0.05, **p<0.01 Student’s t-test. (c,f) Log2 of the fold change of genes activated by Dex in 3134 cells transfected with negative control (si-Neg.C.) or Rev-erbβ / RORβ and RORγ siRNAs. (d,g) Transcriptional changes (log2 fold change of specific knock-down /negative control (Neg.C. siRNA)) of GR upregulated genes (GR-up) in Rev-erbβ (d) and RORβ and RORγ (g) siRNA transfected cells treated with vehicle (-Dex) or Dex. P-values are indicated, Wilcoxon test. (h) A model for gene regulation by GR, ROR and Rev-erb. Both, Rev-erbβ and RORβ/γ, negative and positive GR co-regulators, can bind a composite GR binding site harbouring a GRE.

