## Supplemental data for "Gene activation and repression by the glucocorticoid receptor are mediated by sequestering Ep300 and two modes of chromatin binding"

**Supplementary Figure 1. The transcriptional response to hormone- activated GR includes gene activation and gene repression**. (a) Variation in Pol-II ChIP-seq signal 1 hour after Dex application, presented as $log_2$ of the fold change (FC). The mean Pol-II signal (before and after hormone application) is presented on the horizontal axis in a log2 scale. The red points mark induced and repressed genes, i.e. genes with adjusted z-score > |2|. Black horizontal line indicates the 0.5 cutoff. The black points show the remaining genes. (b) Analysis by RT-qPCR of nascent transcript levels of GR-regulated genes. Samples were collected before and after Dex treatment. Results shown are the average of three independent experiments, with standard deviation presented as error bars.

**Supplementary Figure 2. Stable chromosomal topology of GR-responsive genes.** Spatial domains of the GR-activated genes *Cpan12*, *B3galnt2*, and *Hras*, as well as GR-repressed genes *Ccl2*, *Cxcl5*, and *Il27* in vehicle (EtOH) and Dex (100nM, 1h) treated cells. DHS-seq, GR and Ep300 ChIP-seq, from vehicle and hormone treated cells (100nM, 1h) are shown. Genomic mm9 coordinates; green bar indicates the 4C spatial domain, GR-activated and repressed genes are marked in red and blue, respectively.

**Supplementary Figure 3. STAT6 motifs in 4C domains.** Recognition motifs of STAT6 at DHS and GBS (ChIP-seq) (data from hormone treated cells (100nM Dex, 1h)) within 4C domains of *Ccl2*, *Cited*, *Myc* and *Zfp36l1* (depicted in blue). Genomic mm9 coordinates.

**Supplementary Figure 4. Klf4 in domains of GR activated genes.** (a) Recognition motifs of Klf4 at DHS and GBS within 4C domains of *Tgm2*, and *Bcl2l1* (depicted in red). Genomic mm9 coordinates. (b, c) Levels of Klf4 transcript in 3134 cells transfected with scrambled (si-Neg.C.) or Klf4 siRNA, treated with vehicle (EtOH) or Dex, measured by RT-qPCR (B), and Illumina sequencing (C); cpm=counts per million. (d) RNA levels of Klf4 responsive genes in Klf4 siRNA transfected cells treated with vehicle (EtOH) or Dex. (e) Genome-wide expression patterns from RNA-seq data in Klf4 knock down cells. Variation in RNA levels between two conditions (indicated on the graph title) is presented as $log_2$ of the fold change between the two conditions ($log_2FC$). The mean RNA levels (cpm from 6 samples, three biological repeats for each condition) is presented on the horizontal axis in a $log_2$ scale. The red and blue points mark induced and repressed genes, respectively, i.e. genes with $log_2$ fold change greater than |0.5| and FDR < 0.05. Black horizontal line indicates the |0.5| cutoff. The black points show the remaining genes.

**Supplementary Figure 5. Rev-erb and ROR factors in 3134 cells.** (a) Transcript levels of members of the Rev-erb and ROR families in 3134 measured by q-PCR. Data are presented relative to the levels of GR transcript. RORα, which is low in 3134 cells, was measured also in HepG2 cells as a positive control (shown in white). (b,c,d,e) Levels of RORβ, RORɣ and Rev-erbβ transcript in 3134 cells transfected with scrambled (si-Neg.C.) or gene-specific siRNA, treated with vehicle (EtOH) or Dex, measured by qPCR (b, c) and RNA-seq (d, e). cpm, counts per million. Error bars indicate SD of three biological repeats. * $p<0.05$, ** $p<0.01$ t test. (f, g) Genome-wide expression patterns from RNA-seq data in Rev-erbβ and ROR β+γ knock down cells. Variation in RNA levels between two conditions (indicated on graph title) is presented as $\log_2$ of the fold change between the two conditions ($\log_2$FC). The mean RNA levels (cpm from five samples, two biological repeats for control and two for si) is presented on the horizontal axis in a $\log_2$ scale. The red and blue points mark induced and repressed genes, respectively, i.e. genes with $\log_2$ fold change greater than |0.5| and FDR < 0.05. Black horizontal line indicates the |0.5| cutoff. The black points show the remaining genes.

Supplementary Figure1

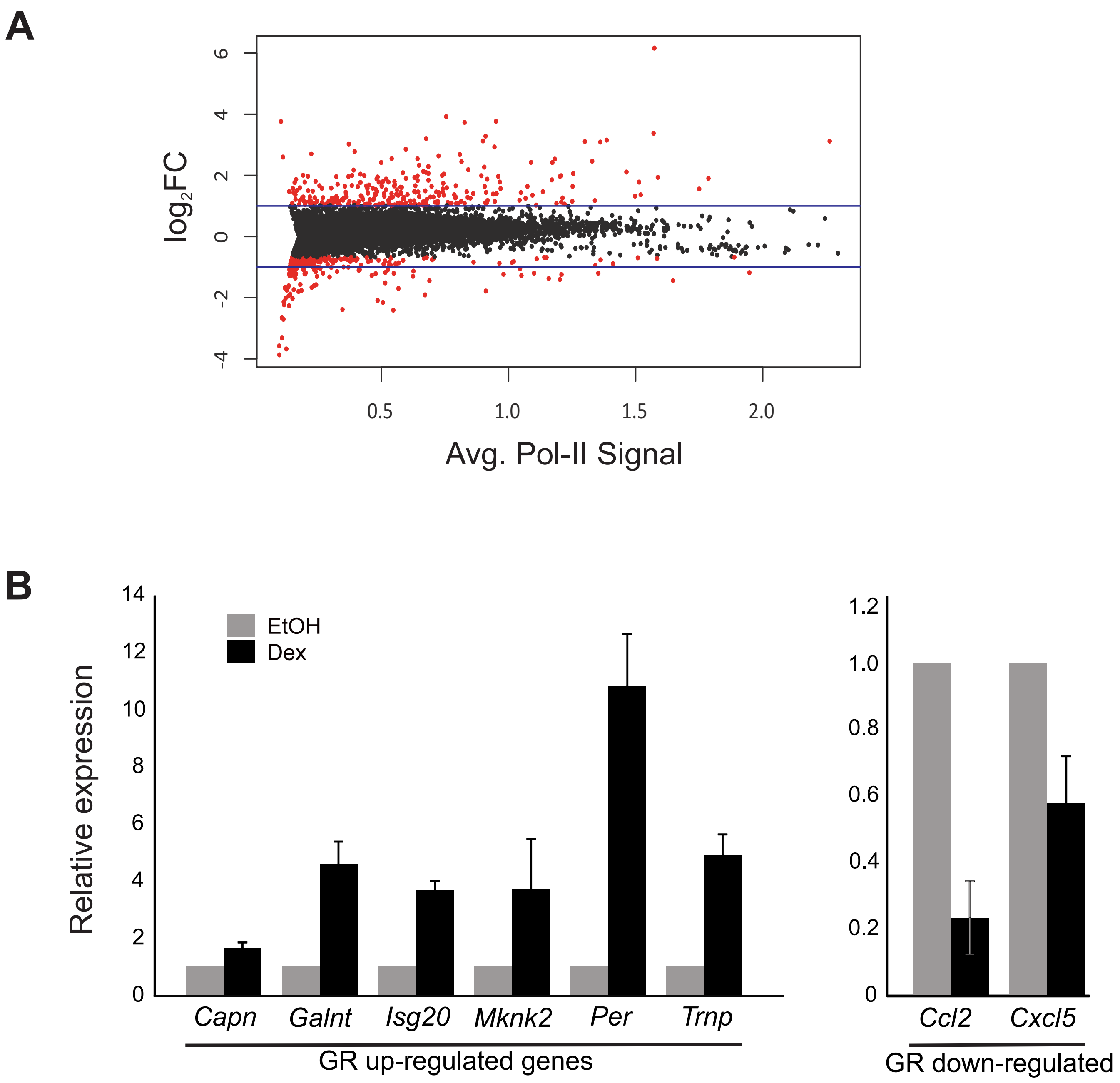

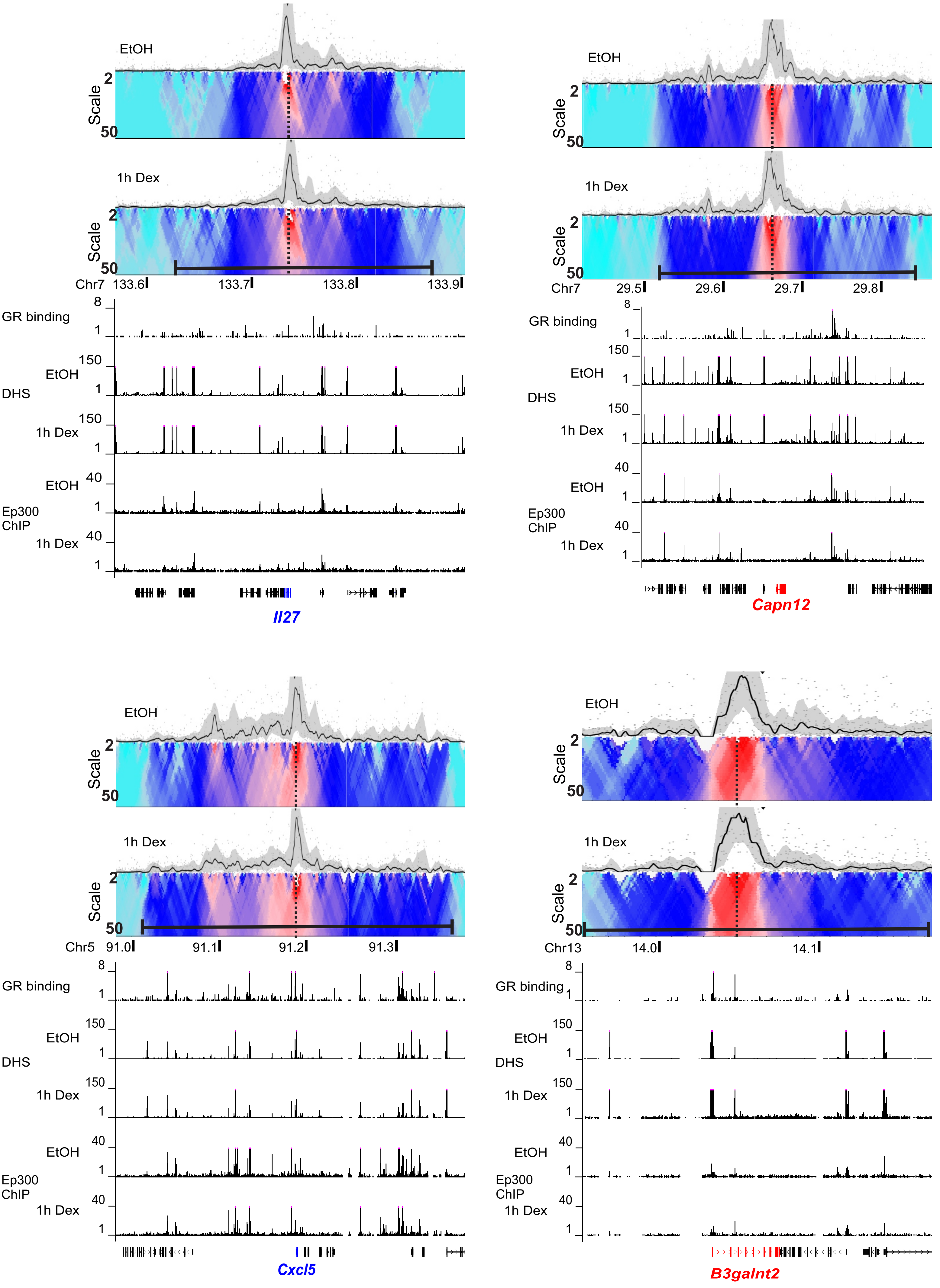


Supplementary Figure2-1

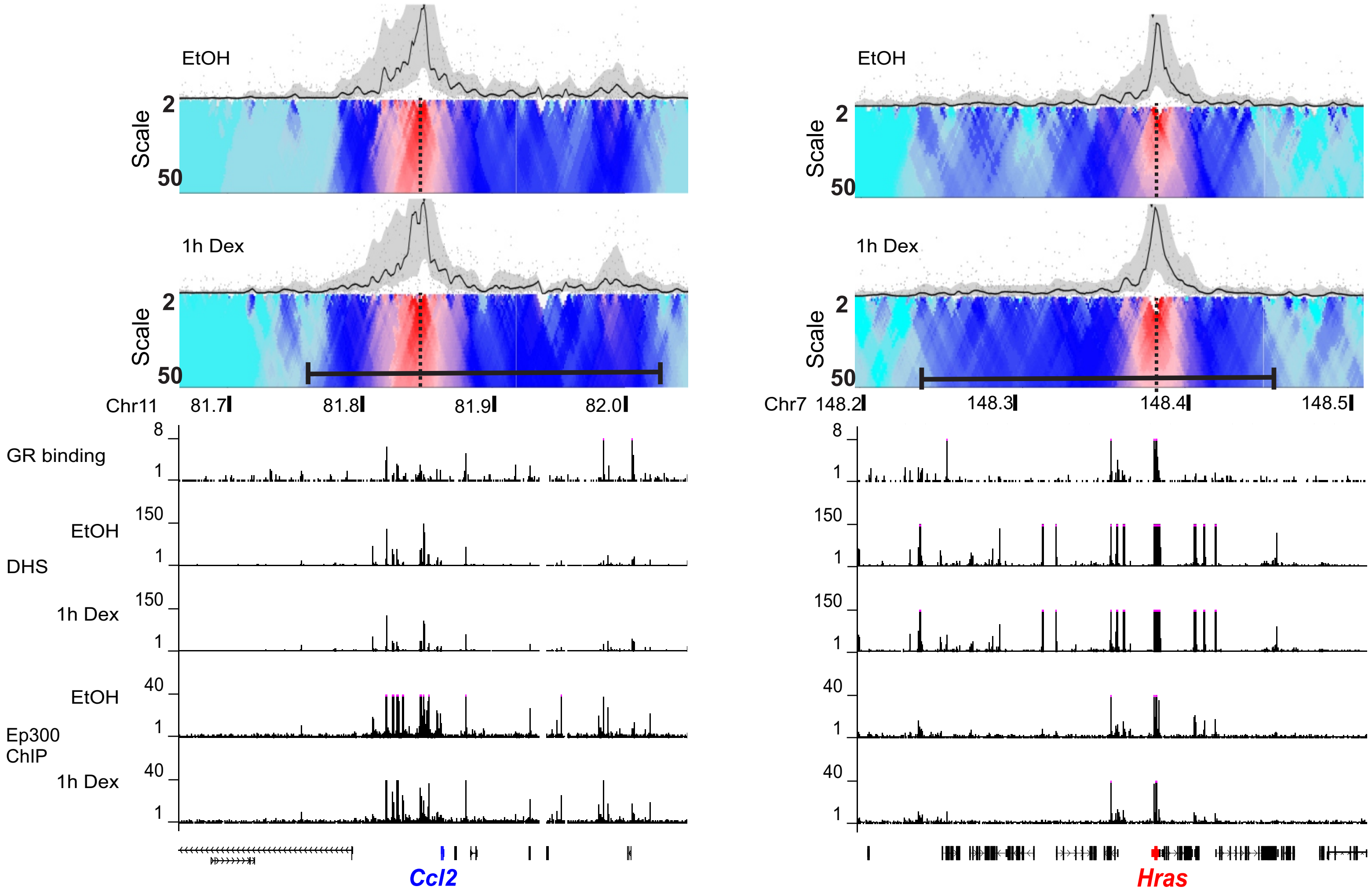


Supplementary Figure 2-2

### Supplementary Figure 3

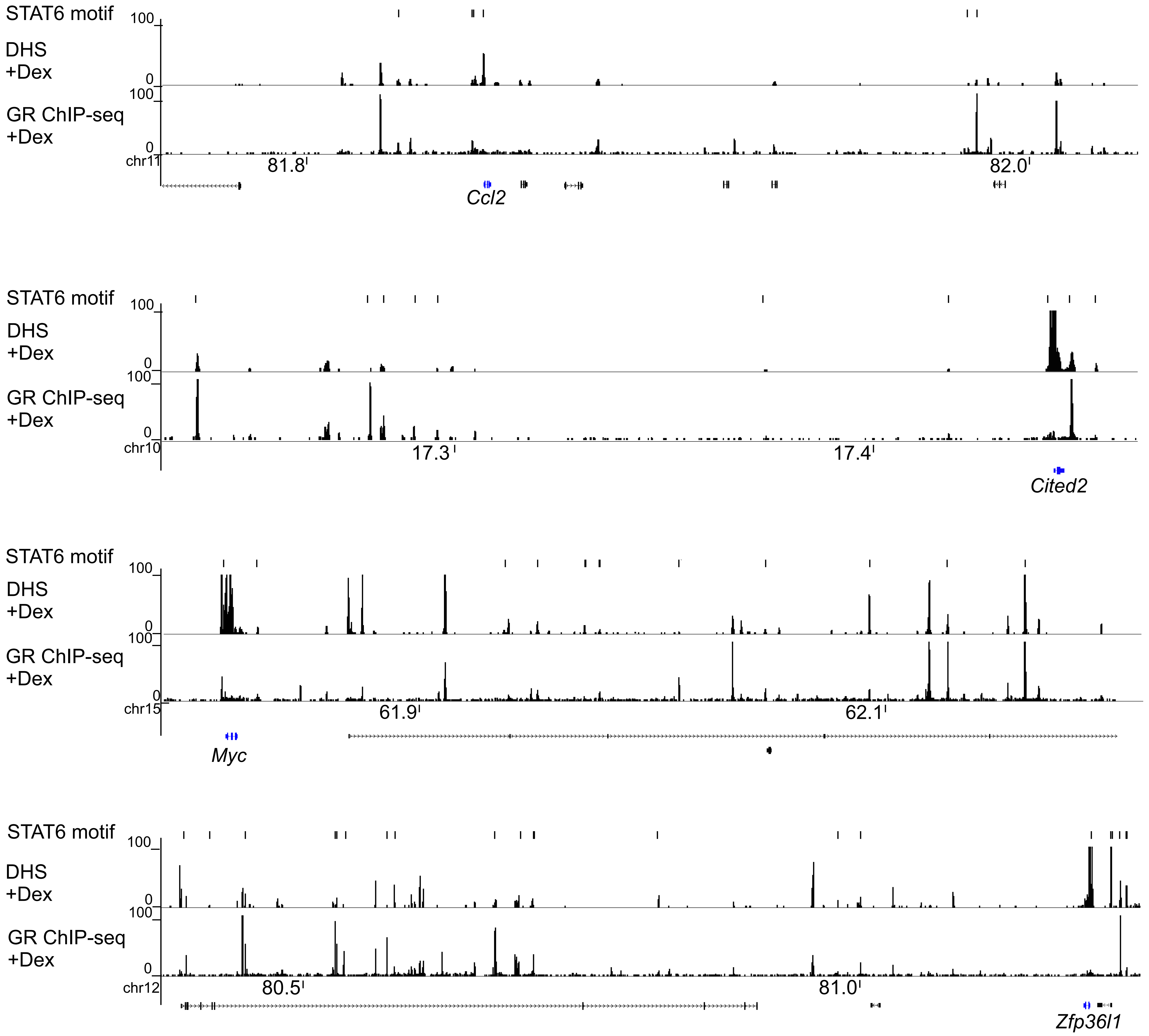

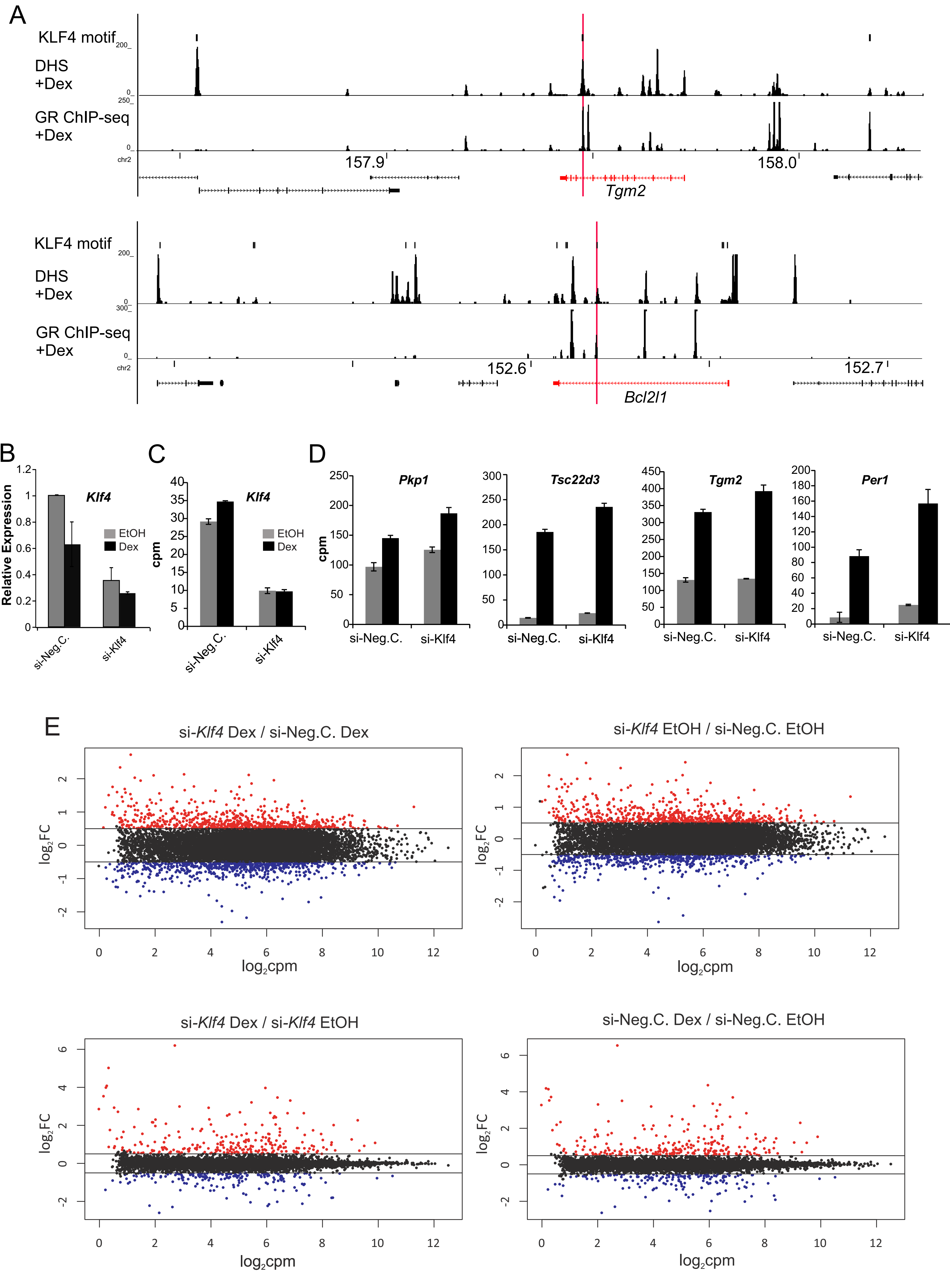


Supplementary Figure 4

### Supplementary Figure5 A-E

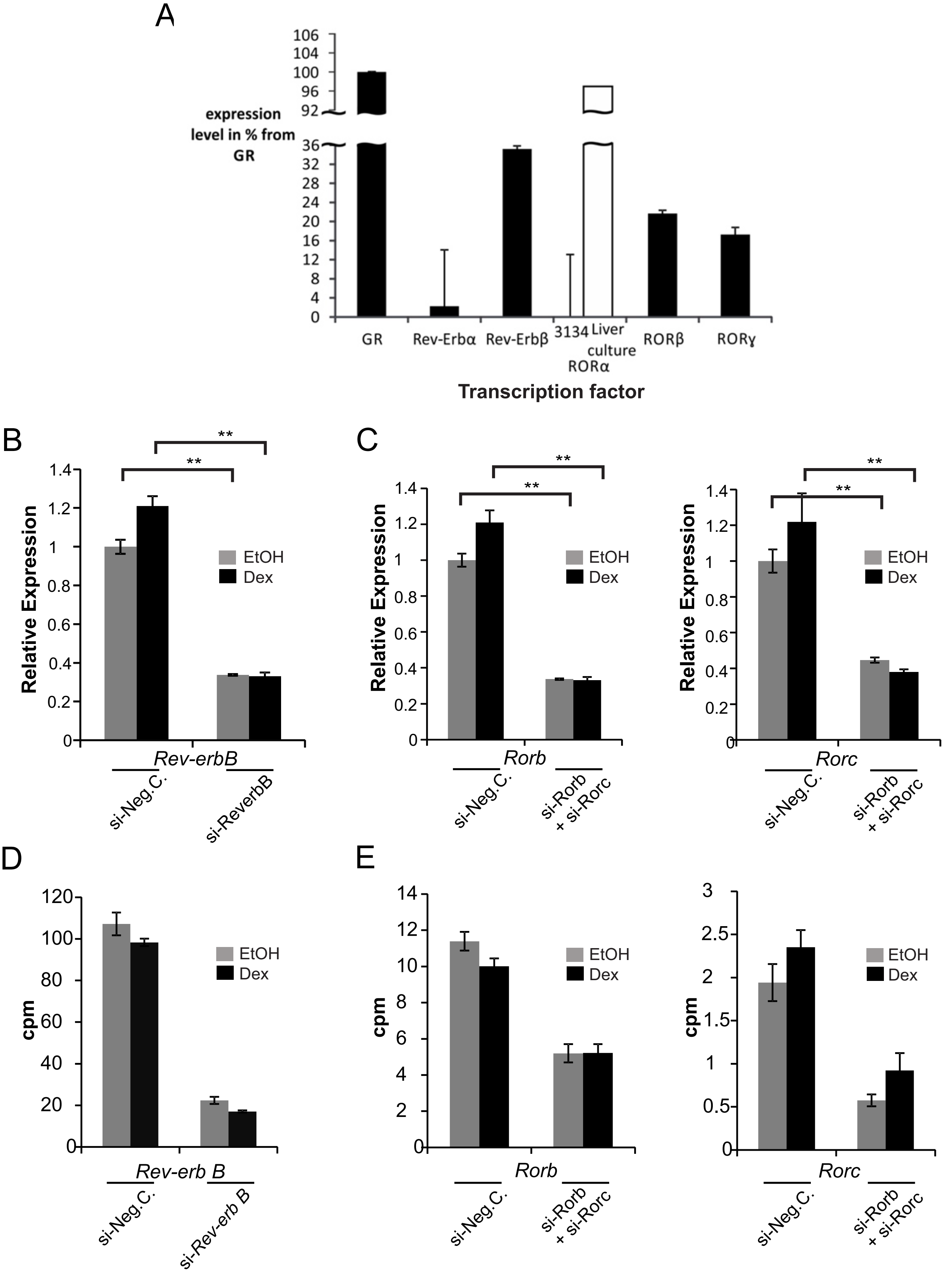

### Supplementary Figure5 F,G

F

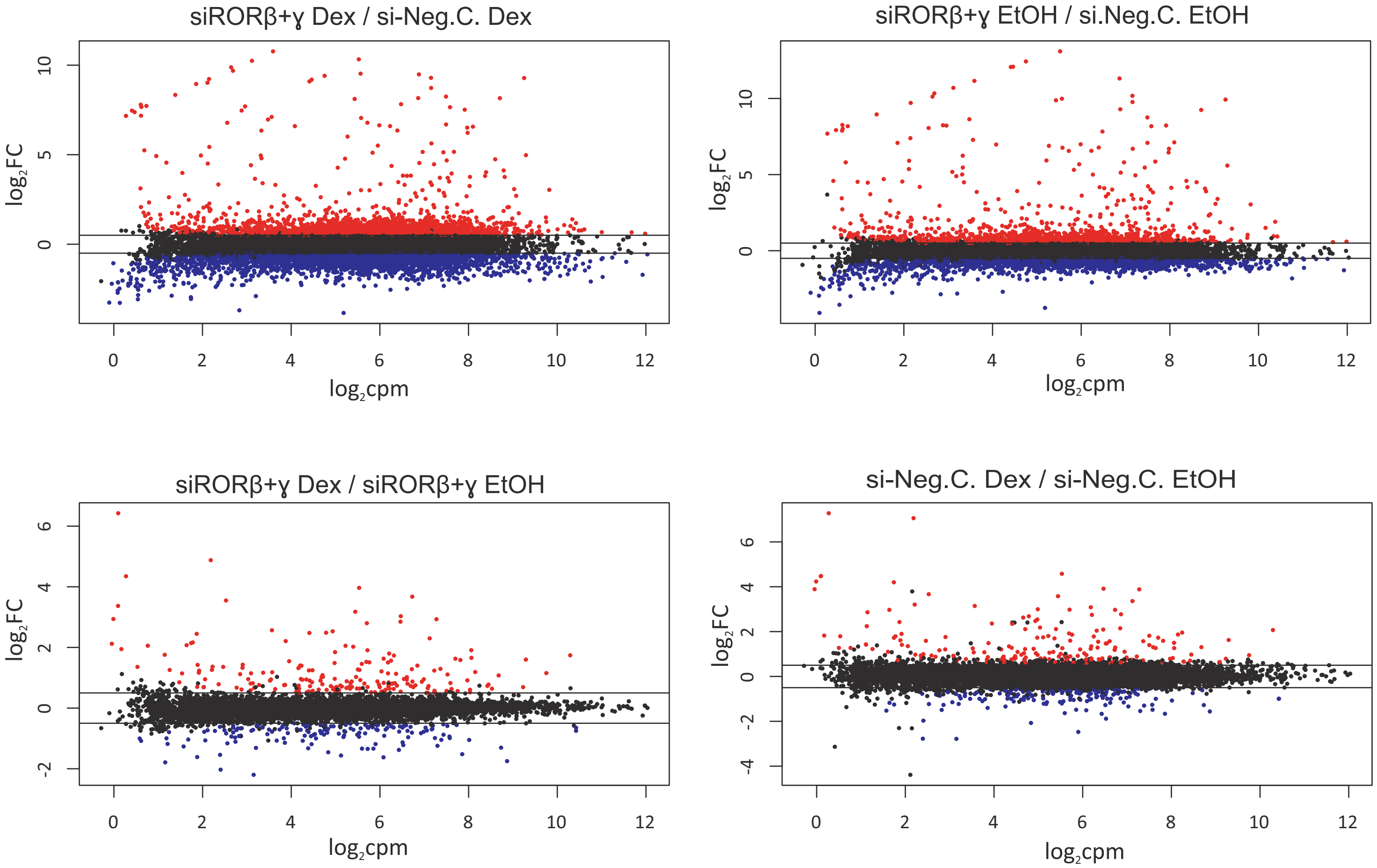


G

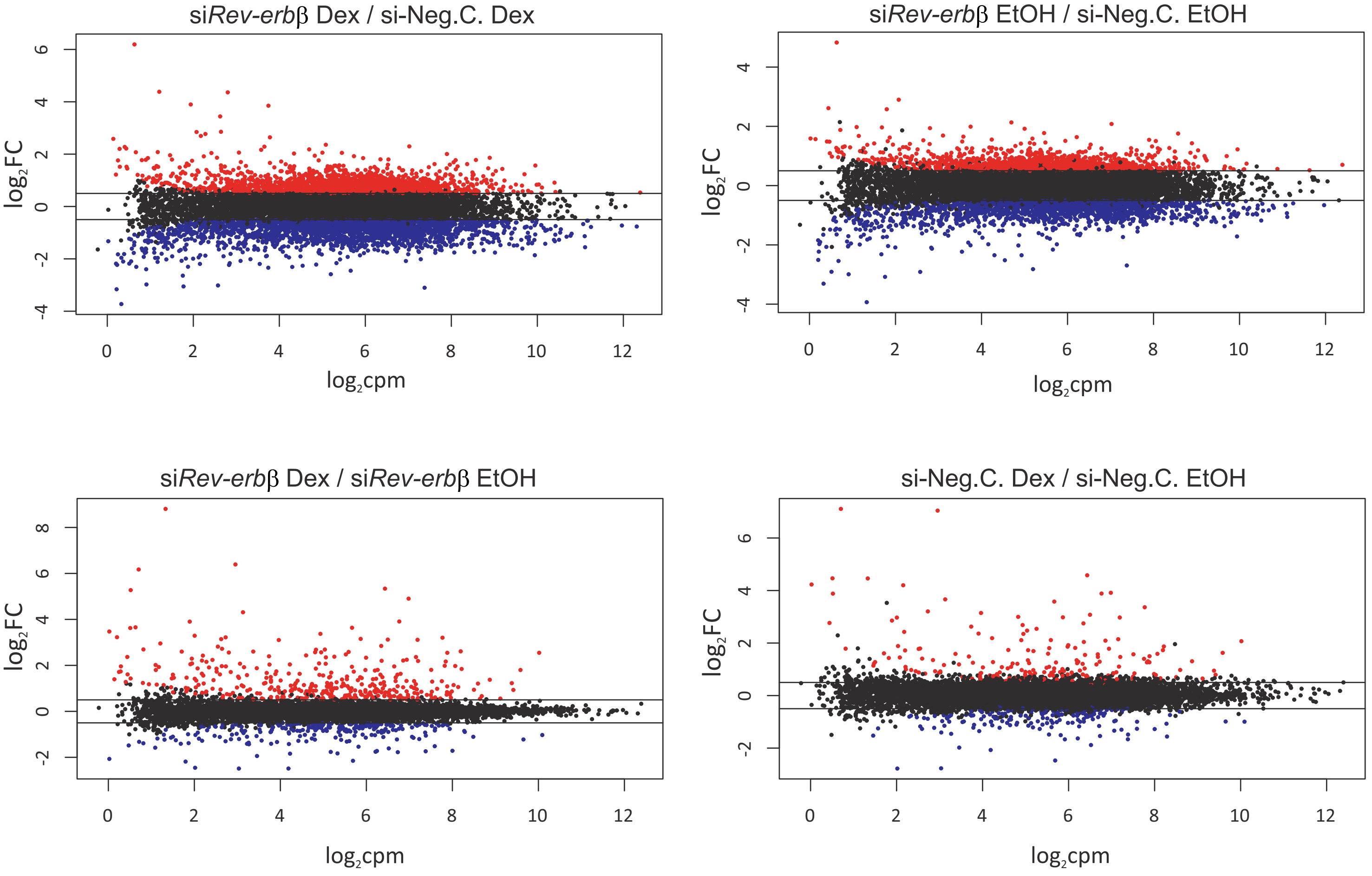
